# SpatialMap: A Scalable Deep Learning Method for Cell Typing in Subcellular Spatial Transcriptomics

**DOI:** 10.1101/2025.07.03.662904

**Authors:** Pengzhen Jia, Yahui Long, Yichao Zhao, Ning Zhang, Zhaoyu Fang, Siqi Chen, Min Li

## Abstract

Subcellular imaging transcriptomics has offered unprecedented resolution of spatial mapping of gene expressions at subcellular level. However, accurate and robust cell type annotation at cellular level remains a challenge. Here we present SpatialMap, a deep weakly supervised cell typing framework that combines graph neural networks (GNN) with transfer learning to annotate cellular identities in subcellular spatial transcriptomics (ST). SpatialMap consistently outperforms state-of-the-art annotation methods on both simulated and real-world ST datasets. It demonstrates strong scalability across diverse tissue types and platforms, including NanoString CosMx, MERFISH and 10x Genomics Xenium. When applied to a mouse primary motor cortex dataset, SpatialMap successfully identified Micro, L4/5 IT, and L6 IT Car3 absent from the reference data, showcasing its capability to discover novel cell types. Furthermore, SpatialMap accurately annotated malignant cells in human liver cancer samples. Its design also enables seamless extension to cross-omics cell typing and cross-platform annotation within proteomics.

## Introduction

The advancements in spatial transcriptomics (ST) technologies have revolutionized biological research by enabling gene expression profiles within tissues while preserving spatial context^1–5^. Recent breakthroughs in subcellular imaging transcriptomics provide unprecedented resolution to elucidate complex tissues at molecular level, and the most popular methodologies include NanoString CosMx^6^, MERFISH^7^, and 10x Genomics Xenium^8^. Despite the breakthroughs of subcellular resolution, new computational methods are still required to infer single-cell information^9^.

Cell type annotation is fundamental for characterizing cellular heterogeneity and understanding tissue biology at single-cell resolution^10^. Traditional annotation methods often rely on manual labelling using marker genes and expert knowledge, which are, however, labor-intensive and time-consuming. Computational methods have been developed to automatically predict cell types, typically leveraging well-annotated single-cell data as reference. Existing computational annotation methods can be roughly divided into non-spatial and spatial methods. Non-spatial methods mainly focus on predicting cell types for single-cell transcriptomics data, including Seurat^11^, Scmap^12^, SingleR^13^, scANVI^14^, CellTypist^15^, and TOSICA^16^. Yet, these methods often face challenges in identifying rare or hard-to-isolate cell types due to the lack of consideration of spatial information.

Spatial methods can be categorized into deconvolution methods, such as CARD^17^, RCTD^18^, cell2location^19^, CellDART^20^, and non-deconvolution methods, such as STELLAR^21^, Spatial-ID^10^, and TACCO^22^. The deconvolution methods, primarily designed for multicellular-resolution spatial transcriptomics platforms like 10x Visium^23^, HDST^24^, and Slide-seq^25^, aim to infer the proportions of different cell types within each spot. While effective at resolving cell type composition, they lack the resolution to assign a unique cell type to individual cell, thereby limiting their ability to reveal fine-grained cellular functions and biological processes. By contrast, non-deconvolution methods assign cell types directly to cells. For example, STELLAR utilizes geometric deep learning to transfer annotation across ST datasets from different regions, tissues, and donors^21^, but its effectiveness is heavily constrained by the requirement for well-annotated reference ST datasets. Spatial-ID combines transfer learning and spatial embedding to enable high-throughput cell typing in ST^10^, while TACCO employs an optimal transport framework to probabilistically map unannotated observations to reference profiles^22^. Despite these advancements, these methods often fail to fully exploit spatial context and typically exhibit suboptimal performance in complex tissues. More importantly, most of these methods face significant challenges in identifying novel and rare cell types, which plays crucial roles in tissue function and disease mechanisms. Moreover, due to inherent distributional differences between multicellular-resolution and subcellular-resolution ST data, methods tailored for multicellular ST often exhibit limited scalability and generalizability when applied to subcellular ST datasets, necessitating the development of bespoke computational methods for cell type identification.

Recently, several segmentation-free cell type annotation methods have been proposed to infer cell types for subcellular ST, such as SSAM^26^, Baysor^27^, FICTURE ^28^, and TOPACT^9^. These methods allow cell type assignment without explicitly segmenting individual cells, thereby reducing dependency on potentially error-prone segmentation. However, they operate on fixed-size grids, transcripts, or spatial neighborhoods to infer cellular identities and thus cannot be easily extended to single-cell-level analyses.

To overcome these limitations, we developed SpatialMap, a deep weakly supervised cell typing framework that combines graph neural networks (GNN) with transfer learning to annotate cellular identities in subcellular spatial transcriptomics. SpatialMap is a scalable and robust framework that leverages well-annotated single-cell data, spatial gene expression profiles, and spatial coordinates to enable high-throughput cell type annotation. We demonstrated the superior performance of SpatialMap on both simulated and real-world subcellular ST datasets, consistently outperforming state-of-the-art annotation methods. SpatialMap exhibits remarkable scalability and robustness across various subcellular ST platforms, including NanoString CosMx, MERFISH, and 10x Genomics Xenium. Notably, it is capable of identifying novel cell types. For instance, in the mouse primary motor cortex dataset, SpatialMap successfully discovered novel populations such as Micro, L4/5 IT, and L6 IT Car3, which were absent from the reference data. Furthermore, we evaluated its transfer learning capability across diverse applications. SpatialMap effectively identified malignant cell populations, including human liver cancer cells, and achieved accurate annotation in both cross-omics cell typing and cross-platform annotation tasks. These results highlight SpatialMap’s versatility and potential to serve as a unified cell typing solution across diverse platforms and omics.

## Results

### Overview of SpatialMap

SpatialMap is a deep weakly supervised cell typing framework that combines graph neural networks (GNNs) with transfer learning to predict cellular identities in subcellular spatial transcriptomics. SpatialMap takes spatial gene expression profiles, spatial coordinates, and reference single-cell data (e.g., scRNA-seq) as inputs, and predicts cell type labels for target spatial transcriptomics (ST). SpatialMap consists of two stages (Fig. 1). In the pre-training stage, the model is first trained on reference single-cell data using both reconstruction loss and classification loss (Fig. 1a). This joint optimization allows the model to capture biologically relevant expression patterns and reliably classify the reference data. In the transfer learning stage, the pre-trained model is applied to the target spatial transcriptomics data to generate preliminary cell type labels (Fig. 1b). High-confidence labels are then selectively retained as pseudo-labels through an adaptive cell-type-wise filtering strategy (see “Methods”). These pseudo-labels are treated as supervision to guide the annotation model, which comprises a GNN-based encoder and an annotator, and enables annotation of the entire ST data. Given the gene expression profiles and spatial coordinates, a spatial neighbor graph is constructed using spatial coordinates, capturing the proximity of spatially adjacent cells (see “Methods”). The GNN-based encoder takes the normalized gene expressions and the spatial neighbor graph as inputs to learn latent representations, which are fed into the annotator to annotate spatial transcriptomics. The model is optimized in a weakly supervised manner, leveraging the pseudo-labels as supervision.

**Figure 1:**
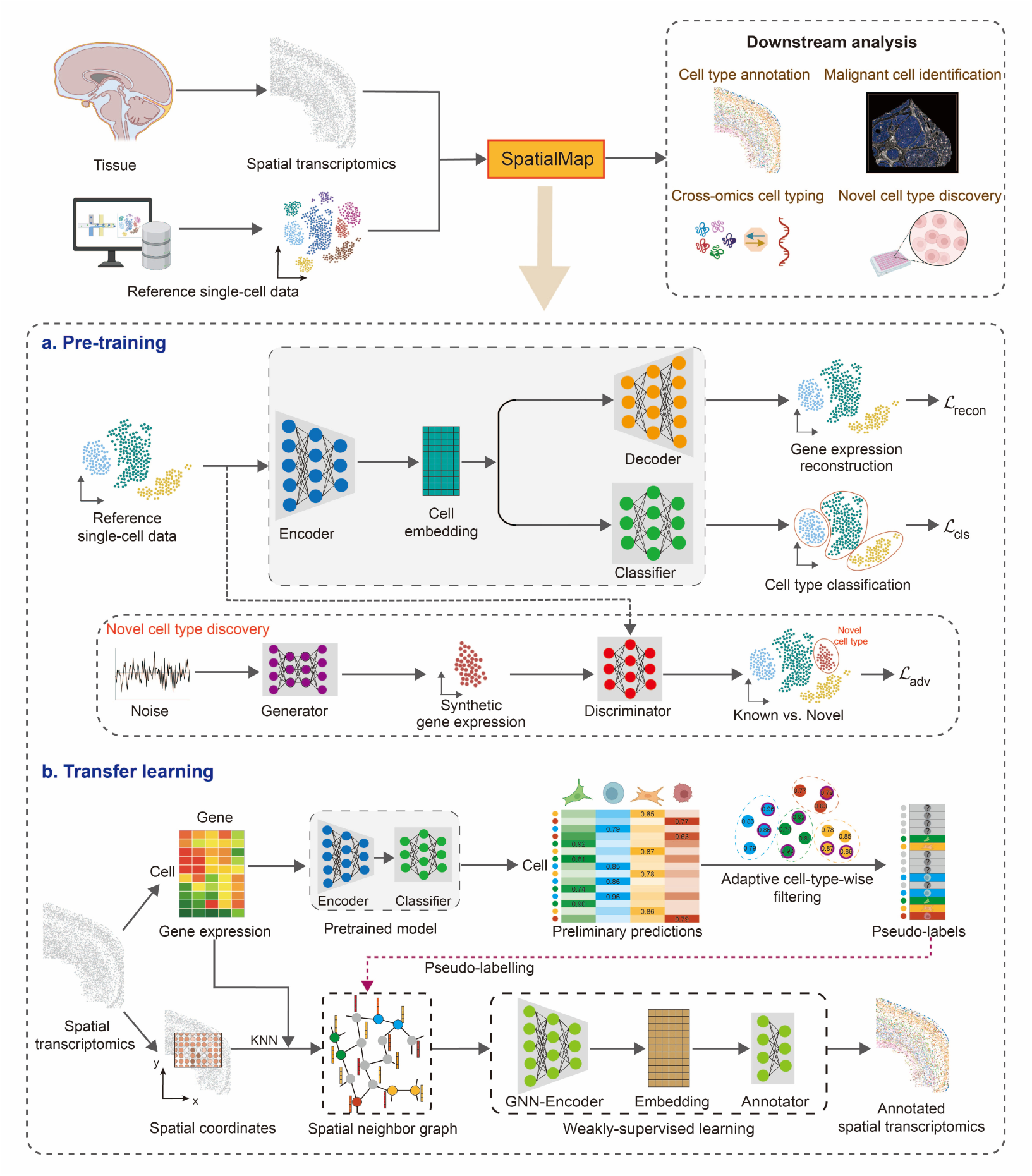
Overview of SpatialMap. SpatialMap is a versatile method for subcellular spatial transcriptomics that enables spatial cell typing, malignant cell identification, cross-omics cell typing, and novel cell type discovery. The SpatialMap framework involves two stages. In the pre-training stage, the model is trained to accurately annotate reference single-cell data using reconstruction loss and classification loss, preserving cell type-specific patterns. A generative adversarial module is incorporated to enable the discovery of novel cell type. In the transfer learning stage, the pre-trained model in the first stage is used for annotating spatial transcriptomics by generating preliminary predictions, followed by high-confidence prediction filtering to obtain pseudo-labels via an adaptive cell-type-wise filtering strategy. Subsequently, the entire spatial transcriptomics are annotated by training a GNN-based annotation framework in a weakly supervised learning manner with pseudo-labels as supervision.

A key point to evaluate a method’s strength is its ability to identify novel cell types present in the target ST dataset, but absent from the reference data. However, it remains a challenge for most existing ST annotation methods. To address this challenge, we incorporate a generative adversarial module into SpatialMap for novel cell type discovery. This module comprises a generator and a discriminator. The generator produces synthetic gene expressions with a random Gaussian noise, representing expression profiles of novel cell types, while the discriminator evaluates whether the input profile originates from a known or a novel cell type. The two components are trained in an adversarial way, allowing the model to robustly detect and characterize previously unseen cell types.

To evaluate the advancement of SpatialMap, we applied it to both simulated and real-world subcellular ST datasets. The simulated dataset was generated from a Stereo-seq mouse brain sagittal section^29^. The real-world ST datasets span human and mouse species and were acquired from various tissue types using different technologies, including NanoString CosMx^6^, MERFISH^30–32^ and 10x Genomics Xenium^8^. SpatialMap consistently outperformed state-of-the-art methods in annotating cell types across six ST datasets. Applied to mouse primary motor cortex, we demonstrated the capability of SpatialMap in uncovering novel cell types. Moreover, SpatialMap exhibited strong versatility for malignant cell identification, cross-omics cell typing, and cross-platform annotation within proteomics.

### SpatialMap accurately annotates cell types in the simulated datasets

We first demonstrated the performance of SpatialMap using simulated spatial transcriptomics datasets. For the simulation, we selected a postnatal day 7 (P7) whole brain sagittal section with a complex spatial pattern, obtained from STOmicsDB^33^. The real dataset contains 96,874 cells with 26,497 measured genes. To generate paired scRNA-seq data, we treated the gene expression profiles from the ST data as scRNA-seq profiles. To mimic the inherent differences in expression profiles between real-world scRNA-seq and ST data, we introduced random Gaussian noise to the ST counts, followed by sampling the ST counts from a Poisson distribution.

For comparison, we benchmark SpatialMap with eight baseline methods, Seurat^11^, cell2location^19^, CellDART^20^, TACCO^22^, spSeudoMap^34^, CIForm^35^, TOSICA^16^, and SPANN^36^. The availability of ground truth allows us to assess the performance with supervised metrics, namely ACC, ARI, NMI, F1-score, AMI, V-measure, and Homogeneity. Overall, SpatialMap achieves more consistent alignment with the ground truth compared to baseline methods (Fig. 2a). Among the methods, Seurat, cell2location, TACCO, CIForm, TOSICA, and SpatialMap produces annotation that are generally closer to the ground truth than those from CellDART, spSeudoMap, and SPANN. Notably, the first six methods successfully recover external granular layer cells with clear spatial patterns. However, only TACCO and SpatialMap are able to recover striatum neuron regions, with SpatialMap exhibiting fewer noises than TACCO. The quantitative results further validate these visual observations (Fig. 2b). To test the robustness of our SpatialMap method, we generated 10 simulated datasets, and evaluated the performance of different methods using the same metrics. The resulting boxplots show that SpatialMap outperforms again the baseline methods with higher median values across seven metrics (Fig. 2c).

**Figure 2:**
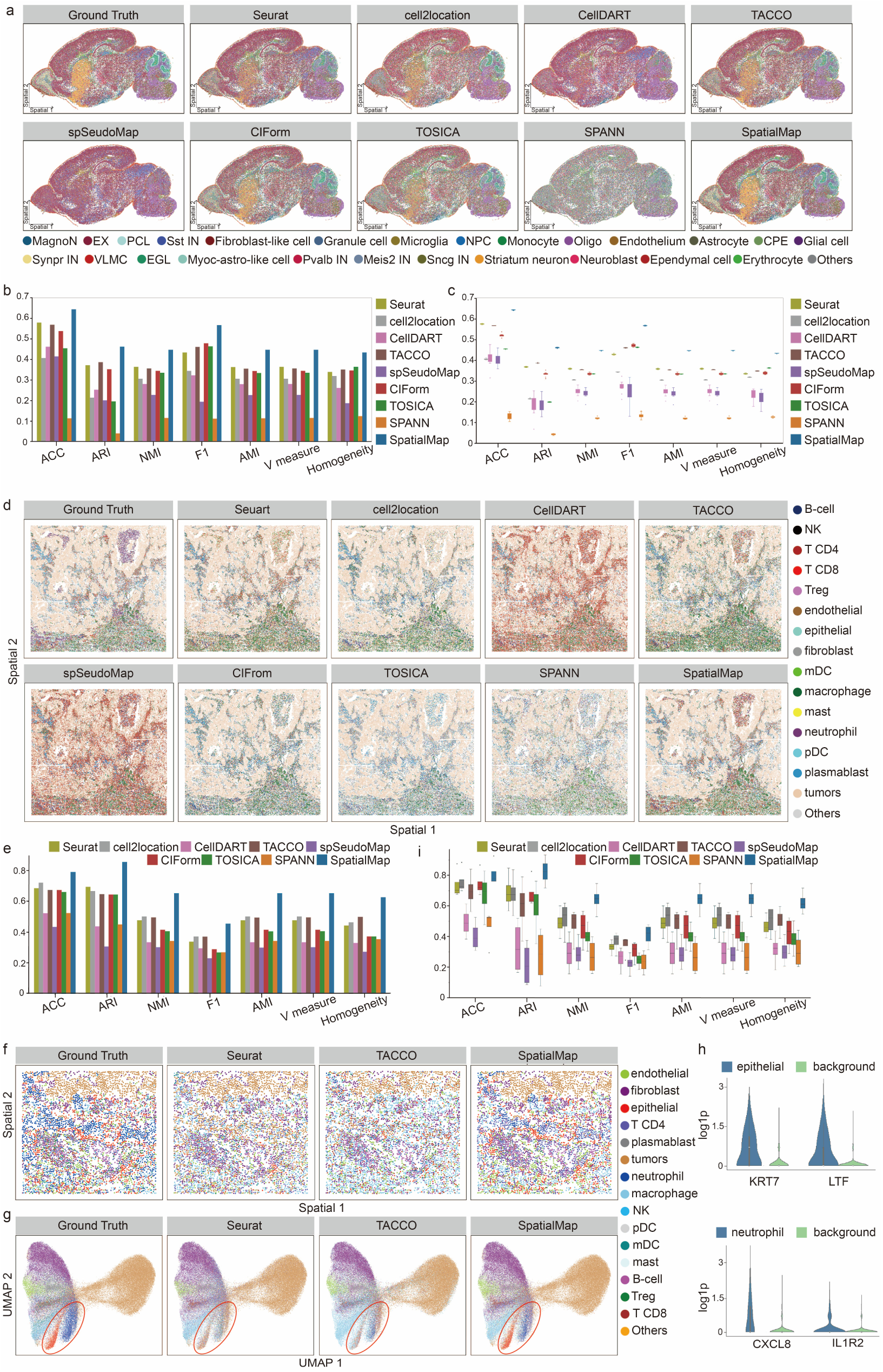
Application to simulated datasets and human NSCLC datasets. **a**. Ground truth and annotation results on a simulated spatial transcriptomics dataset, from SpatialMap and baseline methods, including Seurat, cell2location, CellDART, TACCO, spSeudoMap, CIForm, TOSICA and SPANN. The full names of cell types are: MagnoN: Magnocellular neuron; EX: excitatory neuron; PCL: Purkinje cell; IN: inhibitory neuron; NPC: Neuron progenitor cell; Oligo: Oligodendrocyte; CPE: Choroid plexus epithelium; VLMC: Vascular leptomeningeal cell; EGL: External granular layer cell; Myoc-astro-like cell: Myoc-expressing astrocyte-like cell. **b**, Quantitative comparisons of SpatialMap and baseline methods on the simulated ST dataset using ACC, ARI, NMI, F1, AMI, V measure and Homogeneity. **c**, Boxplots of different methods across 10 simulated datasets. In the boxplot, the centerline, box limits and whiskers denote the median, upper and lower quartiles, and 1.5× interquartile range, respectively. **d**, Ground truth and annotation results on the 9-1 human NSCLC dataset, from SpatialMap and baseline methods, including Seurat, cell2location, CellDART, TACCO, spSeudoMap, CIForm, TOSICA, SPANN and SpatialMap. The full names of cell type are as follows: NK: natural killer cells; Treg: Regulatory T cells; mDC: myeloid dendritic cells; pDC: plasmacytoid dendritic cells. **e**, Quantitative comparisons of SpatialMap and baseline methods on the 9-1 human NSCLC dataset in terms of seven metrics including ACC, ARI, NMI, F1, AMI, V measure and Homogeneity. **f**, Ground truth and spatial distributions of SpatialMap and Seurat, TACCO on the FOV 1, one region from the NSCLC 9-1 sample. **g**, UMAP plots of ground truth and different methods on the FOV 1. **h**, Violin plots of marker gene expressions for epithelial, neutrophil cells predicted by SpatialMap, and background in the NSCLC 9-1 sample. **i**, Boxplots of the seven metrics with scores from 4 human NSCLC datasets. In the boxplot, the centerline, box limits and whiskers denote the median, upper and lower quartiles, and 1.5× interquartile range, respectively.

### SpatialMap accurately annotates cell types in human lung datasets measured by CosMx

Next, we assessed SpatialMap’s performance using real-world ST datasets across various subcellular platforms including NanoString CosMx, MERFISH, and 10x Genomics Xenium. We first applied SpatialMap to a human non-small cell lung cancer (NSCLC) ST dataset^6^ profiled by NanoString CosMx SMI on Formalin-Fixed Paraffin Embedded (FFPE) samples. This dataset was annotated as 16 cell types and one unknown region by original study^6^, namely natural killer (NK) cells, regulatory T cells, myeloid dendritic cells (mDC), plasmacytoid dendritic cells (pDC), fibroblasts, T CD8 cells, plasmablasts, mast cells, T CD4 cells, neutrophils, macrophages, epithelial cells, endothelial cells, B-cells, tumors, monocytes. The reference scRNA-seq dataset^37^ acquired from human lung consists of 52,698 cells and 33,694 genes. The NSCLC ST and reference datasets share 15 cell types and 952 genes.

For evaluation, we compared the performances of SpatialMap and eight competing methods on the Lung-9-1. Visually, SpatialMap detects the spatial distributions of different cell types that agree well with the ground truth. Specifically, the tumor regions identified by Seurat, cell2location, CIForm, TOSICA, and SpatialMap match well with the ground truth while the tumor regions identified by CellDART, TACCO, spSeudoMap, and SPANN mix either T CD8 or other cell types(Fig. 2d, Supplementary Fig. S1). For the non-tumor regions, SpatialMap achieved more accurate annotation than competing methods compared to the ground truth. Notably, only SpatialMap and TACCO are able to identify neutrophils. In contrast, the remaining methods incorrectly maps neutrophils into other regions. Moreover, the spatial distribution of macrophages identified by SpatialMap is significantly more consistent with the annotation. Quantitatively, SpatialMap outperformed competing methods in all metrics (Fig. 2e) with the ARI score of 0.857, which is higher than Seurat (ARI=0.696), cell2location (ARI=0.668), CellDART (ARI=0.439), TACCO (ARI=0.648), spSeudoMap (ARI=0.307), CIForm (ARI=0.646), TOSICA (ARI=0.647), and SPANN (ARI=0.450). The results further validated the superiority of SpatialMap over baseline methods.

Rare cell types often play important roles in key biological process such as immune response. Characterizing rare populations benefits the discovery of novel biomarkers, improving our understanding of cellular heterogeneity. The Lung-9-1 sample consists of 20 FOVs (Field of View). To illustrate the ability of SpatialMap in detecting rare cell types, we additionally visualize FOV 1 (Fig. 2f, Supplementary Fig. S2). The spatial distributions and UMAPs (Fig. 2g, Supplementary Fig. S3) show that compared with the ground truth, SpatialMap is the only method that can accurately isolate epithelial cells and neutrophils from other cell types, while these cell types are misidentified by Seurat, TACCO and other methods. For further validation, we visualize the expressions of the marker genes KRT7, LTF in predicted epithelial cells, and CXCL8, IL1R2 in predicted neutrophils cells, compared to background (Fig. 2h). It shows that these markers are expressed at higher levels in SpatialMap-predicted cell types than in the background.

We further tested SpatialMap and competing methods with the Lung-6, Lung-9-2, Lung-13 samples and measured their performance using the supervised metrics (Fig. 2i). The boxplot shows that SpatialMap outperforms again competing methods in terms of all metrics. The high coherence between predicted cell types and their marker genes across four NSCLS samples underscores the reliability of SpatialMap (Supplementary Fig. S4).

### SpatialMap accurately annotate cell types in mouse brain ST datasets measured by MERFISH

In this second example, we tested SpatialMap on mouse brain datasets measured by MERFISH. We selected three tissue samples from different regions in the brain, including hippocampal formation, isocortex, and hypothalamus regions (Fig. 3a). The mouse hippocampal formation dataset^30^ includes 14 slices with more than 8,000 cells. We collected single-cell transcriptomics data^30^ measured by 10x Chromium Single Cell 3’ v2 as reference. Both datasets are mapped to the Allen whole mouse brain taxonomy and share 500 genes. The visual results of the 36- and 37-th slices show that SpatialMap can accurately recover cell types compared to ground truth (Fig. 3b). We compared SpatialMap with eight competing methods on 14 slices and evaluated their performance using seven supervised metrics. SpatialMap consistently outperforms competing methods in all metrics (Fig. 3c). Specifically, SpatialMap reaches the highest median ACC score of 95.8% with the smallest standard deviation of 0.65%. We also validated the performance of SpatialMap through confusion matrix comparing the predicted cell types with the real cell types (Fig. 3d). SpatialMap accurately identified almost all cell types except CNU-MGE GABA.

**Figure 3:**
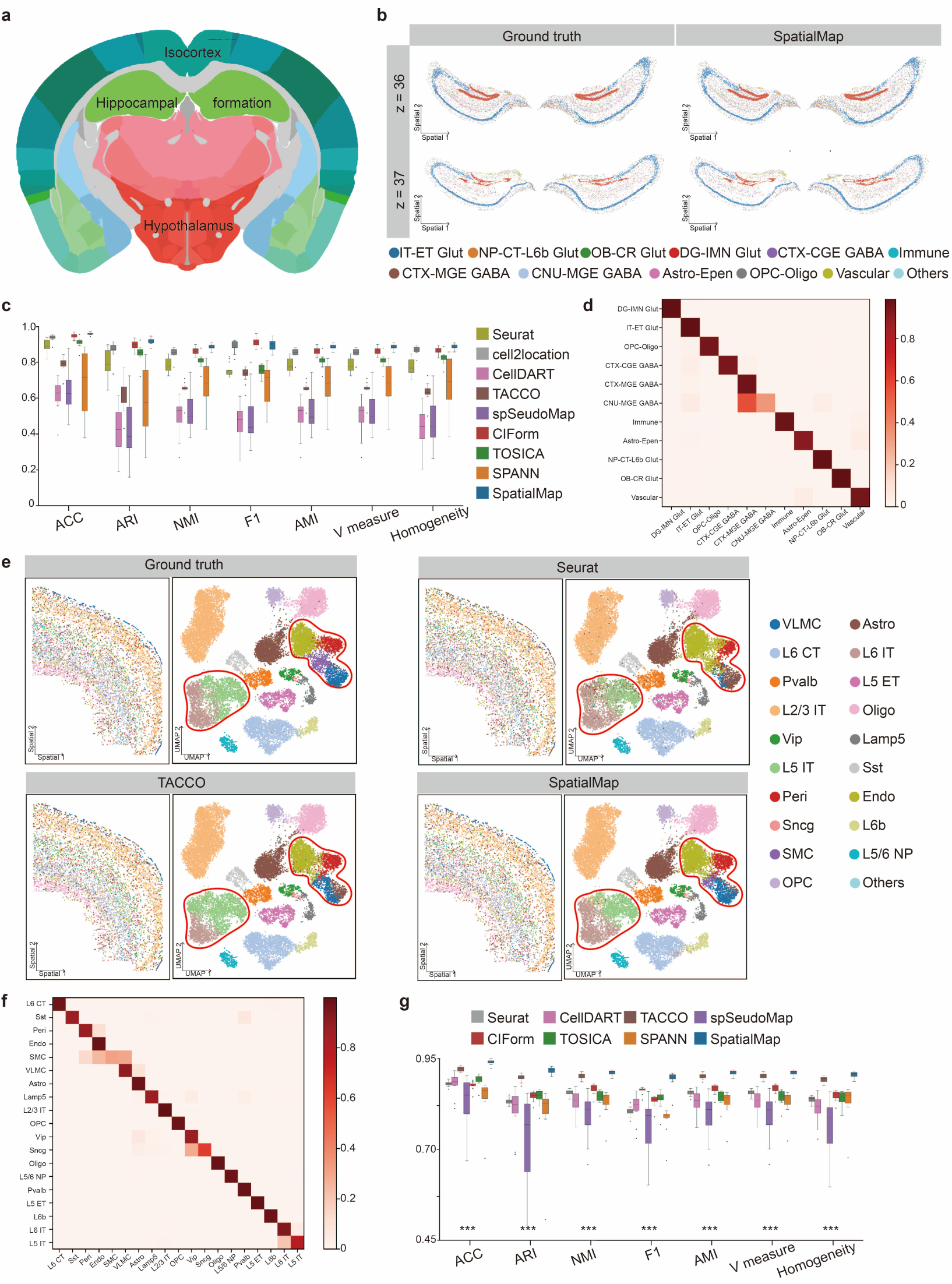
Application to mouse brain datasets. **a**, Allen Brain Altas. **b**, Spatial plots of ground truth and SpatialMap on mouse hippocampal formation datasets. The full names of cell types are as follows: IT: intra-telencephalic; ET: extra-telencephalic; Glut: glutamatergic; NP: near-projecting; CT: corticothalamic; L6b: layer 6b; OB: olfactory bulb; CR: Cajal–Retzius; DG: dentate gyrus; IMN: immature neurons; CTX: cerebral cortex; CGE: caudal ganglionic eminence; GABA: GABAergic; MGE: medial ganglionic eminence; CNU: cerebral nuclei; Astro: astrocyte; Epen: ependymal; OPC: oligodendrocyte precursor cells; Oligo: oligodendrocytes; **c**, Boxplot comparisons of seven metrics for SpatialMap and eight methods, including Seurat, cell2location, CellDART, TACCO, spSeudoMap, CIForm, TOSICA, and SPANN, across 14 mouse hippocampal formation datasets. In the boxplot, the centerline, box limits and whiskers denote the median, upper and lower quartiles, and 1.5× interquartile range, respectively. **d**, Confusion matrix evaluating the agreement between SpatialMap’s annotation (horizontal axis) and known cell types (vertical axis) across 14 mouse hippocampal formation datasets. **e**, Spatial and UMAP visualizations of ground truth, Seurat, TACCO and SpatialMap in the mouse1 sample3 MOP dataset. Left: spatial visualizations in mouse1 sample3, right: UMAP visualizations in mouse1 sample3 slice153. The full names of cell types are as follows: VLMC: vascular leptomeningeal cells; Astro: astrocyte cells; CT: corticothalamic; IT: intra-telencephalic; Pvalb: parvalbumin; ET: extra-telencephalic; Oligo: oligodendrocytes; Vip: vasoactive intestinal polypeptide; Lamp5: lysosomal-associated membrane protein family member 5; Sst: somatostatin; Peri: pericytes; Endo: endothelial cells; Sncg: synuclein gamma; L6b: layer 6b; SMC: smooth muscle cells; NP: near-projecting; OPC: oligodendrocyte precursor cells. **f**, Confusion matrix evaluating agreement between SpatialMap’s annotation (horizontal axis) and known cell types in 12 MOP datasets. **g**, Boxplot comparisons of seven metrics for SpatialMap and seven methods, including Seurat, CellDART, TACCO, spSeudoMap, CIForm, TOSICA, and SPANN, across 12 MOP datasets. In the boxplot, the centerline, box limits and whiskers denote the median, upper and lower quartiles, and 1.5× interquartile range, respectively. *** denotes SpatialMap’s performance for this metric across 12 MOP datasets, with high significance (p-value < 0.001) versus any other methods.

Furthermore, we used the mouse primary motor cortex (MOP) dataset measured by MERFISH from the isocortex region^32^. We used the datasets from 2 mice, consisting of 12 samples with 161,384 cells and 254 genes. We utilized a matched single-nucleus RNA sequencing (snRNA-seq) dataset^38^ as reference. The reference data was labeled using the same MOP cell taxonomy as the ST dataset. The cell types were first categorized into three classes^32^: excitatory neuronal cells (glutamatergic), inhibitory neuronal cells (GABAergic) and nonneuronal cells, and then further subdivided into 23 classes. The MOP ST dataset and reference dataset shared 19 cell types and 253 genes. Visually, compared with the ground truth, SpatialMap performs better than Seurat and TACCO (Fig. 3e, Supplementary Fig. S5). While all of Seurat, TACCO, and SpatialMap can identify clear cortex layer, the annotation generated by SpaitalMap presents more consistent alignment with ground truth. SpatialMap is also the method that is able to accurately identify vascular leptomeningeal cells (VLMC) region compared to ground truth. The UMAP results indicate that SpatialMap accurately recovered smooth muscle cells (SMC) regions while Seurat and TACCO misidentified SMC regions as either endothelial cells (Endo) regions or VLMC regions. SpaitalMap can also distinguish L5 IT from L6 IT, which are labeled with bias by Seurat and TACCO. The Confusion Matrix results across 12 samples also demonstrated again the high accuracy of SpatialMap (Fig. 3f). We compared SpatialMap with seven competing methods on the 12 MOP samples and measured their performance using seven supervised metrics. SpatialMap outperformed baseline methods with a median ACC of 93.9%, a median ARI of 92.0%, a median NMI of 91.3%, and a median F1 of 89.9% (Fig. 3g). The small standard deviation exhibits SpatialMap’s strong robustness across different samples. Wilcoxon tests conducted on the 12 MOP samples for each assessment metrics (p-value<0.001) also demonstrated SpatialMap’s significance (Fig. 3g).

Next, we validated SpatialMap’s performance on the mouse hypothalamic preoptic dataset^31^ measured by MERFISH. This dataset measured from 30 mice contains over one million cells profiled with a 155-gene panel. For evaluation, we selected a single sample comprising 12 tissue slices (from Bregma 0.26 to −0.29) at 50μm interval. The reference was collected from the hypothalamic preoptic region^31^, including 31,299 cells and 27,998 genes. The ST and reference data share 155 genes and eight cell types, including endothelial cells, excitatory neurons, immature oligodendrocytes, mature oligodendrocytes, ependymal cells, microglia, inhibitory neurons and astrocytes. We examined cell type proportions across the 12 slices (Fig. 4a). The cell type compositions predicted by SpatialMap closely matched the ground truth compared to competing methods. We further examined the Pearson correlation coefficients (PCC) between the predicted and real cell type compositions. The results showed that SpatialMap achieved a highest PCC score of 99.9%, followed by TACCO at 98.7%, Seurat at 97.6%, and CellDART at 94.9% (Fig. 4a). The quantitative comparisons of different methods on 12 slices show that SpatialMap achieved the best performance with a median ACC of 92.0%, ARI of 79.8%, NMI of 80.5% and F1 of 91.4%, followed by Seurat and TACCO (Fig. 4b).

**Figure 4:**
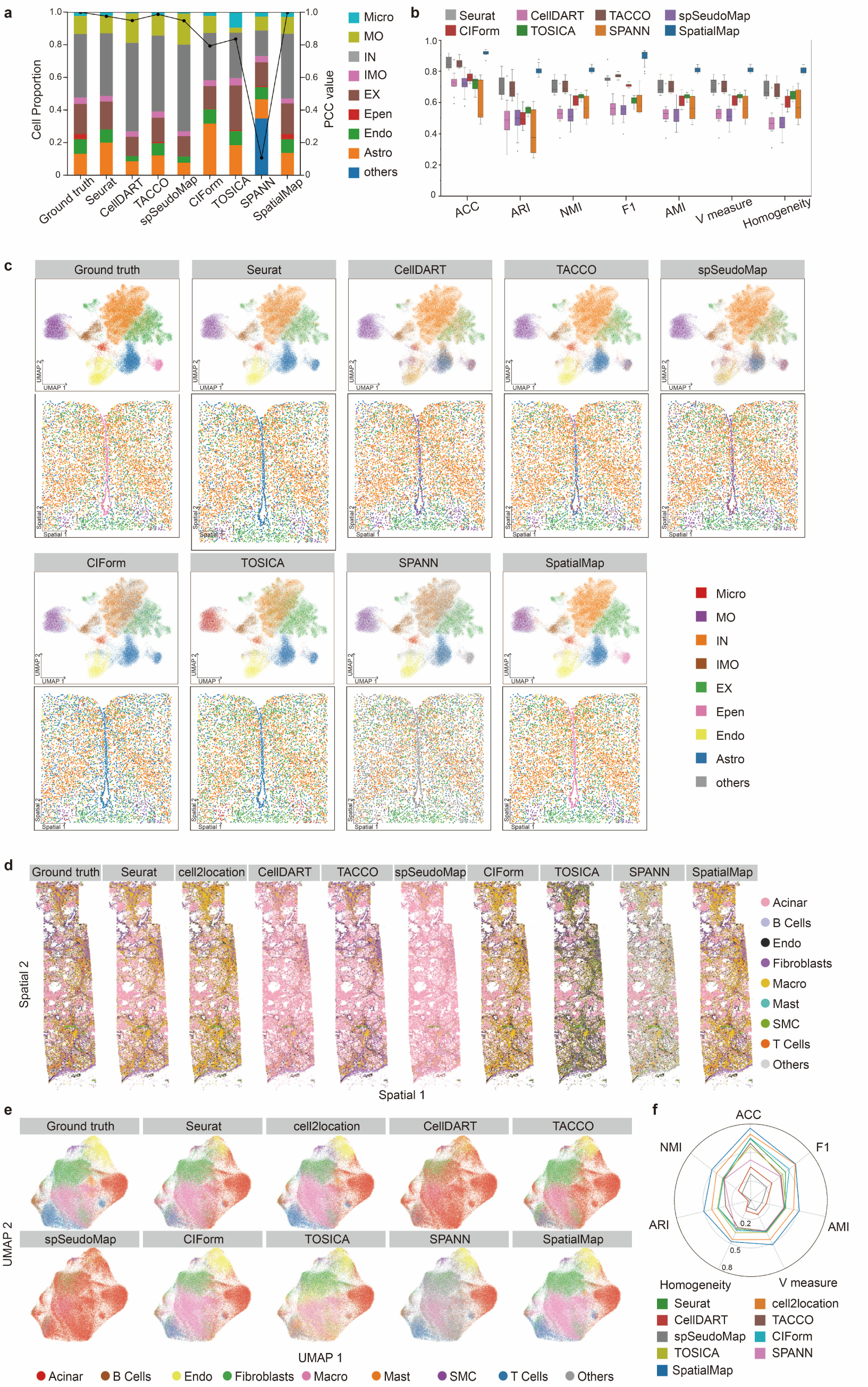
Application to mouse brain and human pancreas datasets. **a**, Cell type proportions (left) and corresponding Pearson correlation coefficients (PCC, right) for the ground truth, SpatialMap, and baseline methods across 12 mouse hypothalamic preoptic region slices. The abbreviations for cell types are explained as follows: Micro: microglia; MO: mature oligodendrocytes; IN: inhibitory neurons; IMO: immature oligodendrocytes; EX: excitatory neurons; Epen: ependymal cells; Endo: endothelial cells; Astro: astrocytes. **b**, Boxplots of the seven metrics for SpatialMap and seven methods across 12 mouse hypothalamic preoptic region slices. In the boxplot, the centerline, box limits and whiskers denote the median, upper and lower quartiles, and 1.5× interquartile range, respectively. **c**, UMAP and spatial plots of the ground truth, SpatialMap, and baseline methods across 12 mouse hypothalamic preoptic region slices and in the representative slice (begma=-0.04). **d**, Spatial plots of the ground truth, SpatialMap, and baseline methods, including Seurat, cell2location, CellDART, TACCO, spSeudoMap, CIForm, TOSICA and SPANN in human pancreatic cancer section. **e**, UMAP plots of the ground truth and different methods. **f**, Radar chart of seven metrics for different methods.

Ependymal cells (Epen) formed a single layer lining the more caudal aspects of the third ventricle^31^ and represent an important rare cell type in mouse hypothalamus regions. Competing methods exhibited low accuracy in identifying Epen, misclassifying them as astrocytes (Astro), mature oligodendrocytes (MO), or other cell types. In contrast, SpatialMap accurately annotated Ependymal cells, further demonstrating its potential in identifying rare cell populations. (Fig. 4c). Besides, given the important roles of inhibitory neurons (IN) and excitatory neurons (EX) in regulating energy balance in the hypothalamus^39,40^, we specifically assessed the performance of various methods in identifying these two cell types (Fig. 4c). Among them, Seurat and SpaitalMap demonstrated superior accuracy in distinguishing IN and EX compared to other competing methods. In contrast, CellDART, TACCO, and spSeudoMap frequently misclassified excitatory neurons as inhibitory neurons, while TOSICA attempted to misclassify inhibitory neurons as excitatory. CIForm and SPANN exhibited greater confusion, mislabeling these neurons as other cell types. Notably, SpatialMap produced annotation with a clearer separation between inhibitory and excitatory neurons than the ground truth. To investigate these discrepancies, we computed the cosine similarity between the conflicting cells and reference expression profiles. The results showed that the cells in question were more closely aligned with SpatialMap’s predictions, highlighting its capability for robust error correction and high classification accuracy (Supplementary Fig. S6). Finally, we validated the spatial distribution of these three cell types (i.e., Epen, IN and EX) identified by SpatialMap using known marker genes^31^ on a representative slice (begma=-0.04, Supplementary Fig. S7). The annotation patterns produced by SpatialMap closely matched the spatial localization of marker gene expressions, further supporting its effectiveness in accurate cell type prediction.

### SpatialMap accurately annotates cell types in human pancreas datasets measured by Xenium

To assess the robustness of SpatialMap across different subcellular ST platforms, we next performed a benchmarking analysis using a human pancreatic cancer dataset measured by 10x Genomics Xenium, which in situ profiled 538 RNA transcripts at subcellular resolution. For annotation, we employed a human pancreas scRNA-seq reference dataset obtained from the DISCO database^41^ as reference. The ST and reference dataset share 474 genes and eight cell types. We benchmarked SpatialMap against eight competing methods, including Seurat, cell2location, CellDART, TACCO, spSeudoMap, CIForm, TOSICA, and SPANN. We first assessed their annotation performance visually (Fig. 4d). Annotation generated by Seurat, cell2location, CIForm, and SpatialMap showed better agreement with the ground truth compared to those generated by CellDART, TACCO, spSeudoMap, TOSICA, and SPANN. This observation was further supported by the UMAP visualization (Fig. 4e). However, Seurat misidentifies endothelial cells as acinar cells, while CIForm and cell2location misidentifies subsets of fibroblasts as macrophages. In contrast, SpatialMap’s annotation demonstrated close alignment with the ground truth across most cell types. Quantitatively, SpatialMap outperformed all baselines, achieving an ACC of 75.5%, an ARI of 50.6%, and an NMI of 51.9% (Fig. 4f). To further explore SpatialMap’s effectiveness in identifying specific cell types, such as acinar cells, we examined the expressions of the acinar marker gene *RNASE1*^42,43^ (Supplementary Fig. S8). SpatialMap’s annotation of acinar cells exhibited strong consistency with RNASE1 expressions, demonstrating its ability to accurately identify hard-to-isolate cell types. Similar consistency was observed across other cell types, confirming the biological interpretability of the annotation based on marker gene distribution.

### SpatialMap enables novel cell type discovery

Novel cell types are often linked to tissue function or dysfunction, yet their identification remains a significant challenge. To date, few existing cell typing methods are capable of reliably detecting such novel populations. To address this limitation, we extended our SpatialMap to support novel cell type identification. For evaluation, we benchmarked SpatialMap against the advanced method SPANN using 12 MOP samples from 2 mice. From the reference dataset, we retained 19 cell types that were shared with the target ST datasets. Additionally, we treated four ST-specific cell types: L4/5 IT, Micro, PVM, and L6 IT Car3, as novel cell types. We first compared both methods on the 50th slice of mouse-1-sample-1. Visually, SpatialMap successfully recovered regions with well-defined boundaries corresponding to the spatial distribution of the novel cell types. In contrast, SPANN incorrectly mislabeled novel cell types outside their expected regions (Fig. 5a), highlighting SpatialMap’s superior ability to uncover novel cell identities.

**Figure 5:**
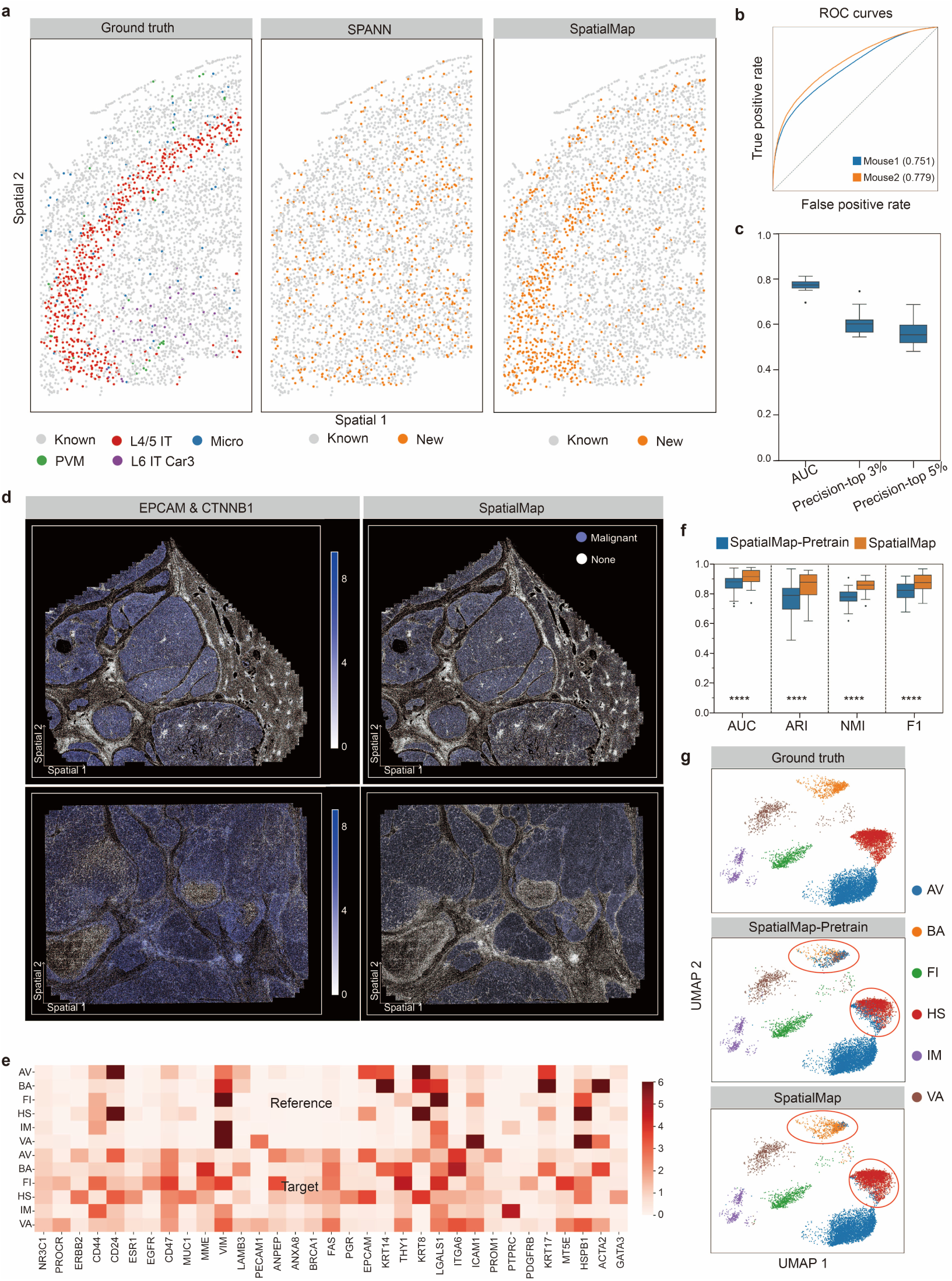
Application to novel cell type discovery, malignant cell identification and cross-omics cell typing. **a**, Spatial visualizations of the ground truth, and predictions by SPANN and SpatialMap in the MOP dataset (mouse1 sample1 slice50). **b**, Quantitative evaluation via AUC of novel cell types predicted by SpatialMap on two mice. **c**, The Boxplots of AUC and precision evaluation across 12 MOP samples. Precision-3% and −5% denote the precision of the top 3% and 5% confident cells identified by SpatialMap. **d**, Spatial visualizations of markers *EPCAM, CTNNB1* and SpatialMap’s malignant annotation in two human liver cancer datasets measured by MERFISH. **e**, Heatmap of the reference (transcriptomic, scRNA-seq) and target (proteomic, CyTOF) profile on 33 matched genes/proteins (x axis) and 6 cell types (y axis) in the human breast atlas datasets. The full names of abbreviate cell types are as follows: AV: alveolar cell; BA: basal cell; FI: fibroblast; HS: hormone-sensing cell; IM: immune cell; VA: vascular lymphatic cell. **f**, Boxplot comparisons of SpatialMap and its invariant SpatialMap-Pretrain across 50 CyTOF samples from 38 patients in terms of AUC, ARI, NMI, and F1. **** denotes SpatialMap’s performance for this metric in 50 CyTOF samples, with very high significance (p-value< 1*e*^-11^) versus SpatialMap-Pretrain. g, UMAP plots of the ground truth, SpatialMap and its invariant SpatialMap-Pretrain in the B1H35-3 sample.

To assess SpatialMap’s capability in identifying novel cell types, we further evaluated the performance of the discriminator, a key component of the model that is responsible for identifying novel cell types. The discriminator outputs probability scores to indicate the likelihood of a cell being a novel cell type. It achieved an AUC score of 75.1% and 77.9% on two mouse datasets, respectively, demonstrating its effectiveness in differentiating novel from known cell populations (Fig. 5b). Besides, across 12 MOP samples, we evaluated its precision on the top 3% and 5% of predicted cells ranked by confidence. The resulting median precision of 60% and 55%, respectively, confirm the discriminator’s utility in prioritizing novel cell candidates (Fig. 5c).

### SpatialMap enables malignant cell identification

Next, we demonstrated the scalability of SpatialMap in identifying malignant cells, a task fundamental to understanding disease progression and treatment response^44,45^. We curated two publicly available human immuno-oncology datasets^46^ generated by MERFISH as target datasets. These datasets, obtained from human liver samples, comprised 568,355 and 598,141 cells, respectively, each profiled with 550 genes. As reference data, we retrieved two human liver scRNA-seq datasets with malignancy annotation from the Tumor Immune Single Cell Hub (TISCH) database^47^. The reference dataset contained 9,597 cells and 438 genes that overlapped with those in the target datasets. Given that the target dataset lacks explicit cell type annotation, we instead assessed SpatialMap’s performance using known malignant cell marker genes, including *EPCAM*^48,49^ and *CTNNB1*^50,51^ (Fig. 5d). Visually, SpatialMap accurately identified malignant regions that showed strong agreement with the expression patterns of these marker genes, underscoring its potential to detect malignant cell populations in complex tumor microenvironments.

### SpatialMap enables cross-omics cell typing

SpatialMap demonstrated strong performance in annotating subcellular spatially transcriptomics samples using scRNA-seq or snRNA-seq data. Beyond spatial omics, we further extended the application of SpatialMap to single-cell proteomics annotation, showcasing its versatility in two cross-modality scenarios: transcriptome-to-proteome and proteome-to-proteome. In the first scenario, transcriptome-to-proteome, we evaluated SpatialMap’s ability to annotate single-cell proteomics data using a human breast atlas dataset^52^, which contains paired CyTOF target data and scRNA-seq reference data. The CyTOF dataset comprises 50 samples from 38 patients and contains 751,970 cells, while the scRNA-seq reference includes 52,681 cells. Both datasets share six common cell types. To align the modalities, we mapped proteins to genes using the UniProt database^53^ and retained 33 matched protein-gene pairs^54^.

A heatmap of expression profiles revealed substantial differences between the CyTOF and scRNA-seq datasets (Fig. 5e). Despite these differences, SpatialMap achieved high annotation accuracy on representative sample (e.g., B1H35-3), showing strong alignment with ground truth labels. Interestingly, SpatialMap achieved more accurate annotation than its variant SpatialMap-Pretrain, which uses only the pre-trained stage, thereby highlighting the critical contribution of the transfer learning stage (Fig. 5f). Moreover, SpatialMap’s annotation displayed clear cell type-specific expression patterns (Fig. 5g). Quantitative evaluation across all 50 samples indicated that SpatialMap achieved better performance with a median ACC of 91.5%, median ARI of 87.7%, median NMI of 85.9%, and median F1 of 87.4%, than SpatialMap-Pretrain. Wilcoxon tests confirmed the significance of these improvements, with all p-values smaller than 1*e*^-11^ (Fig. 5f).

In the second scenario, proteome-to-proteome, we evaluated SpatialMap’s performance in cross-platform single-cell proteomics annotation. We curated two single-cell proteomics datasets derived from murine cell lines, generated by N2^55^ and nanoPOTS^56^. Both datasets included three shared cell lines: an epithelial cell line (C10), a macrophage cell line (RAW) and an endothelial cell line (SVEC), with 762 shared proteins. Notably, the two datasets exhibited substantial differences in feature distributions (Supplementary Fig. S9). Despite these discrepancies, SpatialMap achieved high annotation accuracy of 98.4% when using N2 as reference and nanoPOTS as target, and 99.1% in the reverse setting (Supplementary Fig. S10). These results highlight SpatialMap’s strong potential for accurate cross-platform annotation in single-cell proteomics.

## Discussion

The development of subcellular spatial transcriptomics is revolutionizing molecular biology by providing gene expression profiles at unprecedented molecular resolution. However, despite this advance, inferring accurate single-cell information remains essential for understanding cellular functions and tissue biology. Accurate cell tying is a fundamental step in deciphering cell heterogeneity. Yet, technical limitations such as low capture efficiency pose significant challenges to reliable cell type annotation. In this study, we presented SpatialMap, a deep weakly supervised cell typing framework that combines graph neural networks with transfer learning to accurately annotate cellular identities in subcellular spatial transcriptomics. SpatialMap is a two-stage method that first pre-trains the model using reference single-cell data to preserve cell type-specific patterns, then transfers the pre-trained knowledge into annotating ST. In the first stage, SpatialMap pretrains the model using both gene expression reconstruction loss and classification loss with known cell types. In the second stage, SpatialMap uses the pre-trained model to annotate spatial transcriptomics, generating preliminary cell type labels. The preliminary labels are further filtered with an adaptive cell-type-wise filtering strategy to obtain pseudo-labels. Subsequently, SpatialMap uses a GNN-based encoder and an annotator, trained in a weakly-supervised manner with these partial pseudo-labels, to reliably annotate the entire spatial transcriptomics dataset.

We demonstrated the effectiveness of SpatialMap in annotating spatial tissues using both simulated and real-world datasets across different tissues and subcellular spatial transcriptomics platforms. When annotating simulated whole brain sagittal section, SpatialMap identified cell types that showed more agreement with ground truth compared to competing methods. When applied to subcellular human lung datasets generated by NanoString CosMx, SpatialMap not only accurately delineated tumor and non-tumor regions, but also produced cell type annotation more consistent with the ground truth than those from existing methods. In mouse brain tissue samples from different regions, SpatialMap successfully recovered clear cortical layers with high quantitative accuracy. These results underscore SpatialMap’s robustness and generalizability across multiple tissue types and technological platforms. Notably, we validated the potential of SpatialMap to identify novel cell types using the MOP datasets, a critical challenge that few existing methods are capable of effectively address. Finally, we demonstrated the scalability of SpatialMap with extensions to malignant cell identification and cross-omics cell typing, further highlighting its versatility and potential for extensive spatial omics analysis.

We designed SpatialMap to be user-friendly and flexible, enabling accurate annotation of data generated from various subcellular spatial transcriptomics platforms. SpatialMap has been successfully validated on both simulated Stereo-seq and real-world CosMx, MERFISH and Xenium platforms. We anticipate the advent of subcellular-resolution spatial multi-omics technologies, and we plan to extend our SpatialMap to accommodate these next-generation datasets. SpatialMap is also designed to be computationally efficient in handling challenges from large datasets. The largest dataset we tested contained over 100,000 cells from human pancreas tissue, and it required only 2 min of wall-clock time on a server with Intel(R) Xeon(R) Silver 4114 CPU and GeForce RTX 2080Ti GPU. We believe that SpatialMap represents a powerful and versatile tool for analyzing both current and future subcellular spatial transcriptomics data, advancing downstream biological discovery.

## Methods

### Data Preprocessing

SpatialMap takes gene expression profiles from the reference scRNA-seq (or snRNA-seq) and target ST data as inputs. Common genes between scRNA-seq and ST are first selected. Both reference and target datasets are pre-processed in the same way before model input. Specifically, raw gene expressions are transformed to have zero mean and unit variance for each cell. This cell-wise Z-score normalization helps reduce scale differences across cells and facilitates model optimization. All datasets used in the experiments follow this standardized preprocessing pipeline.

### Graph construction for spatial transcriptomics

Spatial coordinates capture the contextual organization of cells within tissues, and have been demonstrated that it can be exploited to identify similar cell states that are spatially co-located and thus demarcate tissue substructures^57^. Here we use a graph G = (V, E) to characterize cellular neighborhood relationships using spatial coordinates. In the graph G, V ∈ ℝ^*N_cell_*^ represents the set of cells while 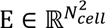 is the set of connected edges between cells with *N_cell_* denoting the number of cells. We calculate the Euclidean distance between centre cell and neighboring cells using the spatial coordinates. For each cell, its neighbors are defined as the top k–nearest cells according to the Euclidean distance (by default, k is set to 3). A ∈ ℝ^*N_cell_×N_cell_*^ is defined as the adjacency matrix of the graph. If cell j ∈ V is the neighbor of cell i ∈ V, *a*_ij_ = 1, otherwise 0.

### Supervised pre-training for initializing annotation model

For cell typing, a deep pre-training framework is first proposed to learn cell type-specific patterns with reference data (e.g., scRNA-seq) as inputs. Let *X*_r_ denote the normalized gene expression matrix of reference data. The encoder consists of stacked fully connected layers, and ReLU activation layers (nonlinear activation layers). It takes as inputs the expression matrix *X_r_* and output latent low-dimensional representation *H_r_* ∈ ℝ^*N_cell_×d*_1_^ with d_1_ denoting representation dimension. The representation 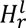 at the l-th layer is formulated as:

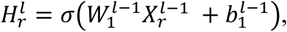

where *W*_1_ and b_1_ denotes trainable weight matrix and bias vector, respectively. *σ*(·) is a nonlinear activation function, i.e., Relu (Rectified Linear Unit).

By contrast, the decoder, sharing a symmetric structure with the encoder, reverses the latent representation *H*_r_ into raw expression space, enabling the latent representation to capture intrinsic gene expression patterns. The reconstructed gene expression *X̃_r_* is formulated as:

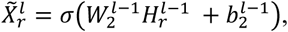

where *W*_2_ and b_2_ denotes trainable weight matrix and bias vector, respectively.

The classifier consists of a linear fully connected layer and a softmax layer. It takes in the latent representation *H*_r_ and output prediction probability matrix *P* ∈ ℝ^*N_cell_×N_type_*^, where *N_type_* denotes the number of ground truth cell types, and *p_ij_* means the possibility of cell i to be assigned into cell type j. Specifically, the classifier is defined as:

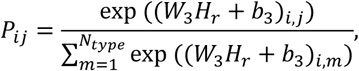

where *W*_3_ and b_3_ denotes trainable weight matrix and bias vector, respectively.

The pre-training model is trained by minimizing the classification loss ℒ_*cls*_ and reconstruction loss ℒ_*recon*_. Briefly, the overall training loss is defined as:

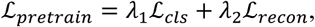

where *λ*_1_ and *λ*_2_ are trade-off factors that control the contributions of the classification and reconstruction losses.

During the experiments, we found that class imbalance degrades classification performance. To address this issue, we used focal loss, an enhanced variant of cross-entropy loss, to train the model, which down-weights well-classified cell types and focuses on difficult and rare cell types. This helps mitigate the bias toward majority cell types and leads to more accurate annotations. Let Y ∈ ℝ*^N_cell_^* denote the ground truth labels. Formally, the classification loss is defined as:

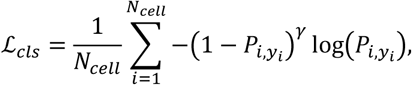

where *γ* is a tunable focusing parameter. *P_i,y_i__* denotes the probability of cell i assigned to ground truth cell type y_i_. The reconstruction loss is formulated as:

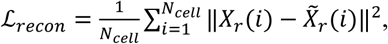

where X_r_ and *X̃_r_* represents the raw and reconstructed gene expressions, respectively.

### Spatially informed transfer learning for annotating spatial transcriptomics

#### Preliminary annotation

After pre-training, we use the classifier trained on the reference data to generate preliminary cell annotations for the target ST dataset, using normalized gene expressions as inputs. However, due to the inherent feature distribution discrepancies between the reference scRNA-seq and ST datasets, these preliminary annotations inevitably include some mislabeled instances. To mitigate the impact of noisy annotation, we introduce a filtering mechanism that retains only high-confidence cell labels. Specifically, the preliminary annotation *T_t_* ∈ ℝ^*N_cell_*^ is first obtained by preserving the highest predicted probability score for each cell 𝑖:

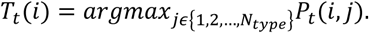

where *P*_t_ ∈ ℝ*^N_cell_^*^×*N_type_*^ is the predicted probability output of the pre-trained model.

To ensure a balanced number of labels across cell types, we implement a cell-type-wise filtering strategy. For each cell type *j*, the top *β* proportion of cells with the highest confidence scores are retained. The confidence threshold is defined as the (1 − *β*)-th percentile of the predicted probabilities within that cell type. The final labels set are treated as pseudo-labels 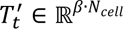, which are defined as:

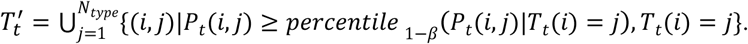

#### Spatially informed ST annotation

With the partial pseudo-labels 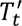, we annotate entire ST sample by fully exploiting the spatial coordinates and gene expression profiles. Assuming that spatially adjacency cells often have similar biological functions or cell identities, a GraphSAGE, an inductive graph representation learning model, is used to encode the spatial graph G to learn cell representations for ST data. Specifically, the latent cell representations *H_t_* ∈ ℝ*^N_cell_×d_2_^* are obtained by iteratively aggregating neighborhood features:

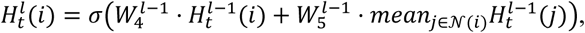

where, 𝒩(*i*) is the set of neighbor cells of the *i*-th cell in *G*. *W*_4_, *W*_5_ are learnable weight matrices. Next, an annotator, consisting of a linear layer and a softmax layer, takes in the latent informative representation *H*_t_ and output predicted probability 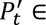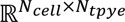.

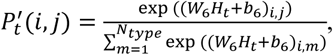

where *W*_6_, b_6_ are trainable weight matrix and bias vector, respectively.

The objective of SpatialMap is to minimize cross-entropy loss in a weakly-supervised manner, measuring the difference between the predicted labels 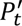 and the known pseudo-labels 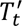.

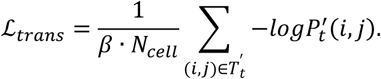

### Novel cell type identification

Novel cell types often represent previously unrecognized cellular functional states that play critical role in advancing our understanding of disease mechanism. However, few existing cell typing methods are able to identify such novel cell types, thereby limiting their applications in real-world scenarios. To overcome this challenge, we extend our SpatialMap to enable the identification of novel cell types that are present in target ST data but absent from the reference dataset.

Building upon the backbone of SpatialMap, we introduce an additional module based on generative adversarial networks (GANs)^58,59^. This module is comprised of a generator and a discriminator. The generator synthesizes artificial gene expression profiles by transforming Gaussian noise into biologically plausible patterns, which are used to represent novel cell types. An adversarial training strategy is employed to optimize the module. The generator is trained to produce synthetic pseudo-gene expression data that resembles real gene expression profiles of known cell types to deceive the discriminator, while the discriminator is simultaneously optimized to accurately distinguish between real expressions from known cell types and synthetic expressions representing novel cell types. The objective of this module is to minimize adversarial loss ℒ_adv_.

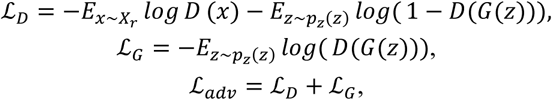

where, *G*(·) and *D*(·) are generator and discriminator, respectively. *z* ∼ *p_z_*(*Z*) denotes the random noise *z* sampled from a prior Gaussian distribution *p*_z_(*Z*). After training, the discriminator is capable of identifying whether a gene expression profile originates from a known or novel cell type, thereby enabling the robust discovery of previously unobserved cell types in spatial transcriptomics data.

### Cross-omics cell typing

SpatialMap is further extended to support cross-omics cell typing for single-cell multi-omics data, including both transcriptome-to-proteome and proteome-to-proteome. For transcriptome-to-proteome cell typing, we utilized matched single-cell multi-omics datasets in the experiments. Prior to model training, 33 matched protein-gene pairs were filtered and selected to serve as model inputs based on the UniProt database. Due to the absence of spatial information in the datasets, an encoder, consisting of fully connected layers with ReLU activation functions, was designed to learn latent cellular representations for each omics modality instead of GNN-based encoder. For proteome-to-proteome cell tying, we first identified and retained the shared proteins between the reference and target datasets. An encoder, similar to the transcriptome-to-proteome scenario, was then used to learn latent representations of proteins instead of GNN-based encoder.

### Implementation details of SpatialMap

For datasets acquired from different technological platforms, the Adam optimizer and learning rate of 0.0001 were used in pre-training and annotation stage. Type-wise filtering strategy with *β* =0.3 was conducted on all the datasets. To account for differences in feature distribution across technological platforms, a tailored weight factor for reconstruction loss in pre-training stage was empirically assigned to each one. The weight factor was 10.0 for CosMx platform and 2.0 for all the other platforms. We also provided a default parameter set that would work for most users on most data types. The training epochs of the pre-training stage and the annotation stage for all the real-world datasets acquired from different technological platforms were 30 and 100, respectively.

### Comparison with baseline methods and evaluation

To showcase the effectiveness of SpatialMap in ST cell type annotation, we compared SpatialMap with eight state-of-the-art methods, the single-cell transcriptomics method Seurat, TOSICA, CIForm and ST methods cell2location, CellDART, TACCO, spSeudoMap and SPANN.

#### Seurat

Seurat is a popular R library for comprehensive single-cell transcriptomics analysis. We followed the pipeline to perform the projection of reference data onto a query object. To be specific, we preprocessed the datasets and then found a set of anchors between the reference and query object with the FindTransferAnchors function. We used the TransferData function to classify the query cells based on the reference data.

#### TOSICA

TOSICA is a multi-head self-attention deep learning model using biologically understandable entities. TOSICA perform cell type annotation with gene sets as masks. For human datasets including simulated datasets, NSCLC datasets and pancreas datasets, we used the provided human_gobp as masks. For mouse datasets, we perform cell typing based on mouse_gobp masks. We followed the usage pipeline and ran TOSICA with default parameters.

#### CIForm

CIForm is a Transformer-based single-cell annotation method, involving gene embedding, positional encoding, Transformer encoder and classification. We followed the tutorial in CIForm’s GitHub for cell typing. For datasets with different genes, we adjust the number of heads in the self-attention mechanism to the value that is divisible by the number of genes and closest to the official setting of 64, to ensure usability.

#### cell2location

cell2location employs a Bayesian model to estimate the spatial distribution of cell types in the ST data of a given tissue using scRNA-seq as reference. For scRNA-seq reference data, we followed the tutorial and performed the filter_genes function with ‘cell_count_cutoff=5’; ‘cell_percentage_cutoff=0.03’ and ‘nonz_mean_cutoff=1.12’. We used the default negative binomial regression model to estimate reference cell type signatures. We used ‘max_epochs=250’; ‘batch_size=2500’ and ‘train_size=1’ to train the model. Next, we found shared genes and perform spatial mapping with hyperparameters: ‘max_epochs=30000’, ’ N_cells_per_location=1’ (for cell type annotation in single-cell resolution) and ‘detection_alpha=20’. Finally, we chose 5% quantile of the posterior distribution, representing the value of cell abundance. It is worth noting that cell2location is based on a strict negative binomial assumption and therefore cannot be used in the MOP and mouse hypothalamic preoptic datasets used in this manuscript.

#### CellDART

CellDART implements an adversarial discriminative domain adaptation to infer the cell fraction in ST datasets. All experiments were implemented using the recommended parameter choices, including ‘num_markers=20’, ‘npseudo=20,000’, ‘nmix=8’, ‘iteration number = 3,000’ and ‘mini-batch size = 512’.

#### TACCO

TACCO is a computational framework for the transfer of annotations to cells and their combinations. TACCO is composed of an optimal transport model extended with different wrappers. Followed the tutorial in mapping single cells into space, we used the tc.tl.annotate function to perform cell type annotation, with ‘multi_center=False’ to use the original annotation categories.

#### spSeudoMap

spSeudoMap is another domain adaptation method to utilizes reference single-cell data to create a cell mixture resembling the spatial data and predicts the spatial cell composition. All experiments were implemented using the default parameters.

#### SPANN

SPANN employs coupled-variational autoencoder to learn latent representations and introduces cell-type-level alignment with prototype optimal transport. We preprocessed datasets, constructed SPANN model and trained the model with the recommended parameters. For the simulated dataset (known types) annotation, we tried to adjust the Resolution parameter (the expected minimum proportion of known cells) to 1 to predict fewer novel types. However, the result (ACC of 31.0%) is not as good as the default ‘Resolution=0.5’ (ACC of 42.5%). Therefore, we ran SPANN at default ‘Resolution=0.5’.

## Data availability

All the public datasets are freely available as follows. The raw Stereo-seq dataset MouseBrain_P7_section1_singlecell.h5ad used for simulation is available from https://db.cngb.org/stomics/datasets/STDS0000139/summary. The real-world datasets are freely available as follows. The human NSCLC ST datasets are available from https://nanostring.com/products/cosmx-spatial-molecular-imager/nsclc-ffpe-dataset/ and the NSCLC scRNA-seq dataset is available from https://gbiomed.kuleuven.be/english/cme/research/laboratories/54213024/scRNAseq-NSCLC. The mouse hippocampal formation ST datasets are available from https://alleninstitute.github.io/abc_atlas_access/descriptions/MERFISH-C57BL6J-638850.html and the hippocampal formation scRNA-seq dataset is available from https://alleninstitute.github.io/abc_atlas_access/descriptions/WMB-10Xv2.html. The mouse MOP datasets are available from https://doi.brainimagelibrary.org/doi/10.35077/g.21 and the reference snRNA-seq 10x v3 B dataset is available from https://assets.nemoarchive.org/dat-ch1nqb7. The mouse hypothalamic preoptic region datasets are available from https://datadryad.org/stash/dataset/doi:10.5061/dryad.8t8s248 and the reference scRNA-seq dataset is available from https://www.ncbi.nlm.nih.gov/geo/query/acc.cgi?acc=GSE113576. The human pancreas dataset is available from https://www.10xgenomics.com/products/xenium-human-pancreatic-dataset-explorer and the reference pancreas scRNA-seq dataset is available from disco_pancreas_v01.h5ad in DISCO database (https://www.immunesinglecell.org/). The human liver ST datasets are available from https://info.vizgen.com/ffpe-showcase and the reference datasets are available from T010016 and T010045 in TISCH database (http://tisch.comp-genomics.org/home/). The scRNA-seq and CyTOF human breast atlas datasets are available from GEO (accession no. GSE180878) and https://data.mendeley.com/datasets/vs8m5gkyfn/1 respectively. The N2 and nanoPOTS murine cell line datasets are available from the MassIVE data repository (accession no. MSV000086809) and the MassIVE data repository (accession no. MSV000084110).

## Code availability

The code is implemented in Python and publicly available at GitHub (https://github.com/pengzhenjia/SpatialMap.git). Comprehensive tutorials are provided to ensure reproducibility and facilitate its application.

## Author Contributions

M.L. and Y.L. conceptualized and supervised the project. P.J. and Y.Z. designed the model. P.J. developed the SpatialMap software. P.J., Y.L., and M.L. wrote the manuscript. P.J., Y.L., Y.Z., Z.F., N.Z., and S.C. performed the data analysis. P.J., Y.L., and M.L. contributed to figure design and generation.

## Acknowledgements

The work was supported by the Fundamental Research Funds for the Central Universities of Central South University.

## Competing Interests

The authors declare no competing interests.

## Notes

### Competing Interest Statement

The authors have declared no competing interest.

