## Supplementary for "SpatialMap: A Scalable Deep Learning Method for Cell Typing in Subcellular Spatial Transcriptomics"

### Supplementary Figures

**Figure S1.** Spatial visualizations of the consistency between the annotation of each method and the ground truth cell malignancy in the 9-1 human NSCLC dataset.
**Figure S2.** Spatial visualization of the FOV 1 region on NSCLC 9-1 sample colored by the ground truth cell types and annotation of methods, including Seurat, cell2location, CellDART, TACCO, spSeudoMap, CIForm, TOSICA, SPANN and SpatialMap.
**Figure S3.** UMAP visualization on NSCLC 9-1 sample colored by the ground truth cell types and annotation of methods, including Seurat, cell2location, CellDART, TACCO, spSeudoMap, CIForm, TOSICA, SPANN and SpatialMap.
**Figure S4.** Heatmap for the average marker gene expression across cell type identified by SpatialMap in 4 human NSCLC datasets. Each row represents a specific cell type annotated by SpatialMap, while each column corresponds to a marker gene. Color indicates log1p-transformed mean expression of each gene within cell type, calculated by first averaging raw expression values across all cells of that type, then applying the log1p transformation.
**Figure S5.** UMAP visualization and spatial visualization of the mouse1 sample3 primary motor cortex dataset colored by the ground truth cell types and annotation of methods, including Seurat, CellDART, TACCO, spSeudoMap, CIForm, TOSICA, SPANN and SpatialMap. a: whole sample3, b: mouse1 sample3 slice153.
**Figure S6.** Cosine similarity between the conflicting cells (SpatialMap vs. Ground truth of inhibitory neurons (IN) and excitatory neurons (EX)) and reference expression profiles.
**Figure S7.** Spatial visualization of the mouse hypothalamic preoptic region dataset (begma=-0.04) for ependymal cells, excitatory neurons and inhibitory neurons annotated by SpatialMap with corresponding marker genes.
**Figure S8.** Spatial visualization of each cell type of SpatialMap's annotation (right) on the human pancreas dataset with corresponding marker gene (left).
**Figure S9.** UMAP embeddings showing the distribution of two proteomics datasets: N2 and nanoPOTS.
**Figure S10.** Quantitative metrics of SpatialMap in Case1 (N2 to nanoPOTS) and Case2 (nanoPOTS to N2) of cross-platform proteomics annotation, including ACC, ARI, NMI, F1.
**Figure S11.** Ablation studies of SpatialMap's on four real-world subcellular datasets. a: human NSCLC measured by CosMx. b: mouse primary motor cortex measured by MERFISH. c: human pancreas measured by Xenium.
**Figure S12.** The influence of different β on SpatialMap's performance on three subcellular technology platform. a: the human NSCLC dataset measured by CosMx. b: the human pancreas dataset measured by Xenium. c: the mouse primary motor cortex dataset measured by MERFISH. SpatialMap adopts β=0.3 for cell type annotation.
**Figure S13.** The influence of different k on SpatialMap's performance on three subcellular technology platform. a: the human NSCLC dataset measured by CosMx. b: the human pancreas dataset measured by Xenium. c: the mouse primary motor cortex dataset measured by MERFISH. SpatialMap shows high robustness to k.

### Supplementary Tables

**Table S1.** Description of all the datasets used in this study.

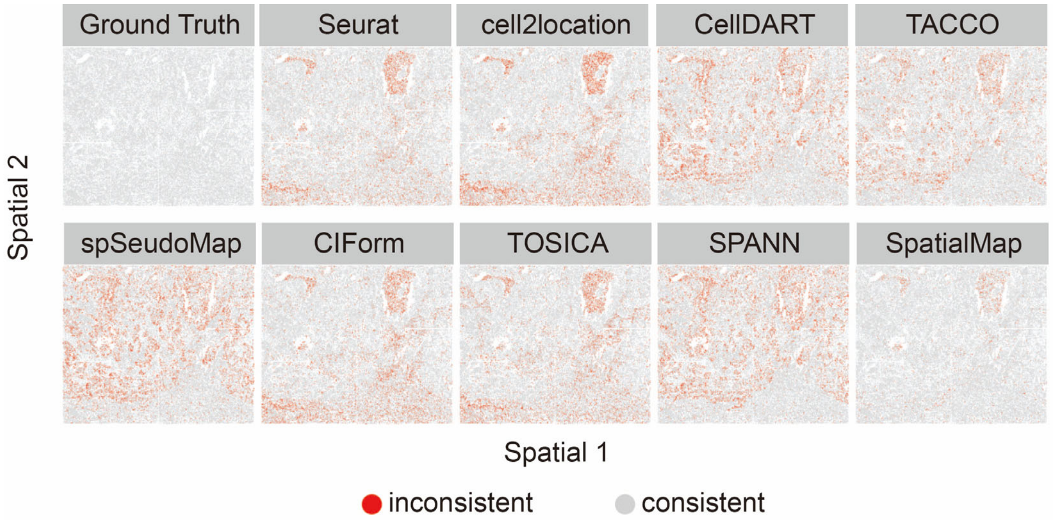


Figure S1. Spatial visualizations of the consistency between the annotation of each method and the ground truth cell malignancy in the 9-1 human NSCLC dataset

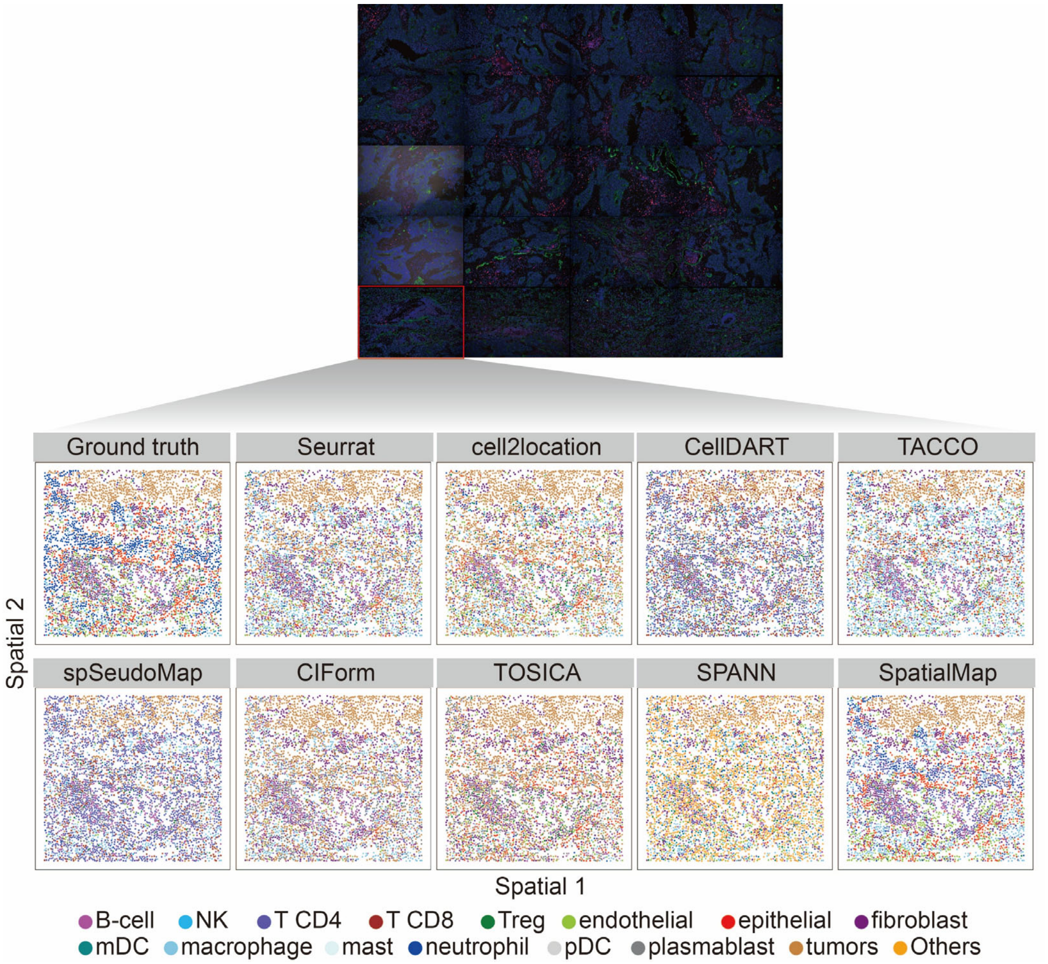


Figure S2. Spatial visualization of the FOV 1 region on NSCLC 9-1 sample colored by the ground truth cell types and annotation of methods, including Seurat, cell2location, CellDART, TACCO, spSeudoMap, CIForm, TOSICA, SPANN and SpatialMap.

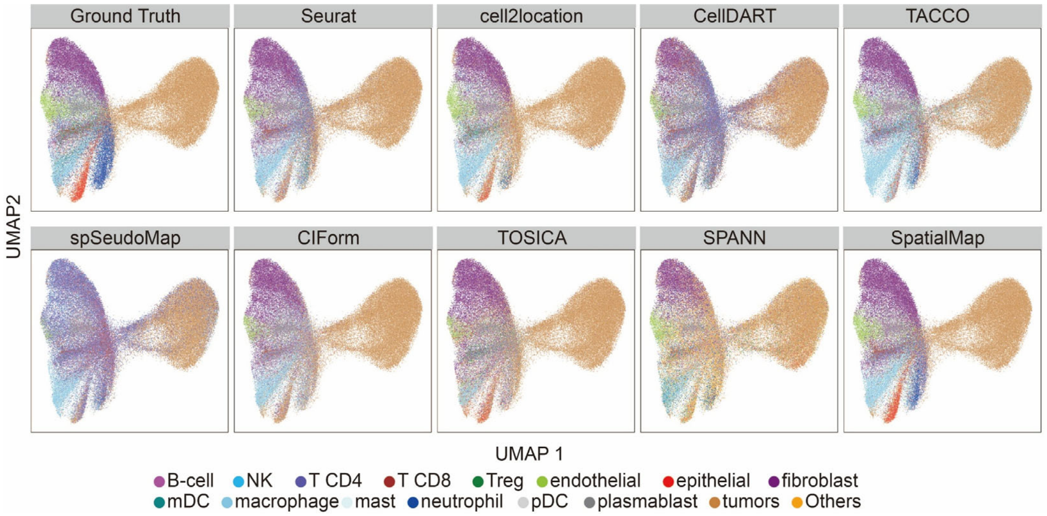


Figure S3. UMAP visualization on NSCLC 9-1 sample colored by the ground truth cell types and annotation of methods, including Seurat, cell2location, CellDART, TACCO, spSeudoMap, CIForm, TOSICA, SPANN and SpatialMap.

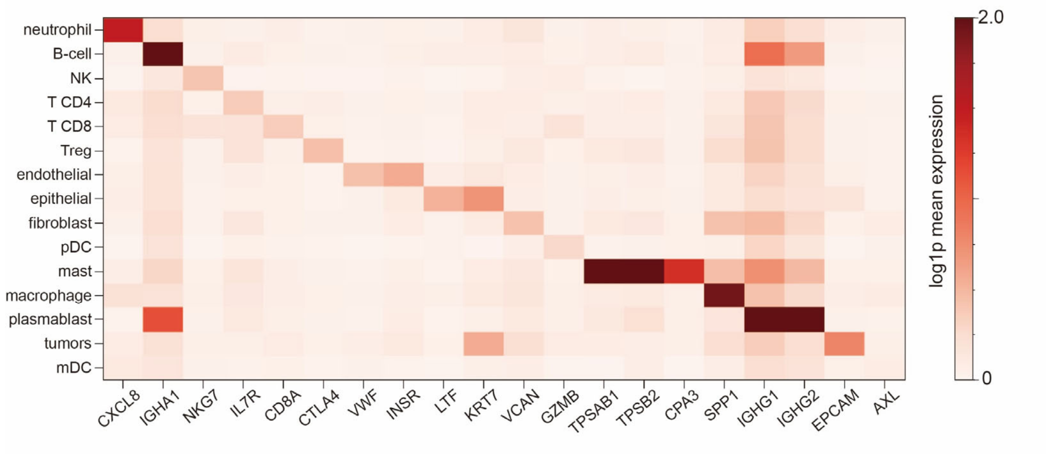


Figure S4. Heatmap for the average marker gene expression across cell type identified by SpatialMap in 4 human NSCLC datasets. Each row represents a specific cell type annotated by SpatialMap, while each column corresponds to a marker gene. Color indicates log1p-transformed mean expression of each gene within cell type, calculated by first averaging raw expression values across all cells of that type, then applying the log1p transformation.

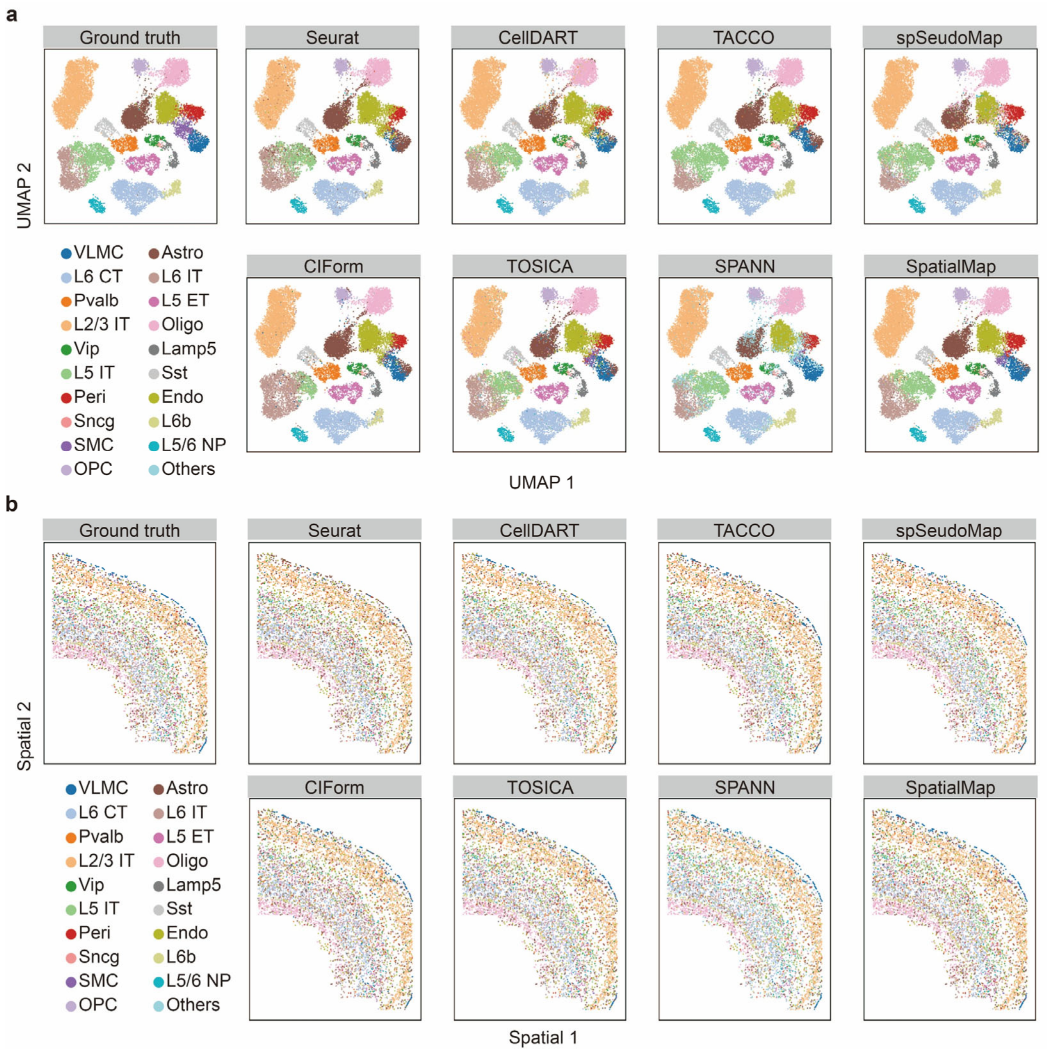


Figure S5. UMAP visualization and spatial visualization of the mouse1 sample3 primary motor cortex dataset colored by the ground truth cell types and annotation of methods, including Seurat, CellDART, TACCO, spSeudoMap, CIForm, TOSICA, SPANN and SpatialMap. a: whole sample3, b: mouse1 sample3 slice153.

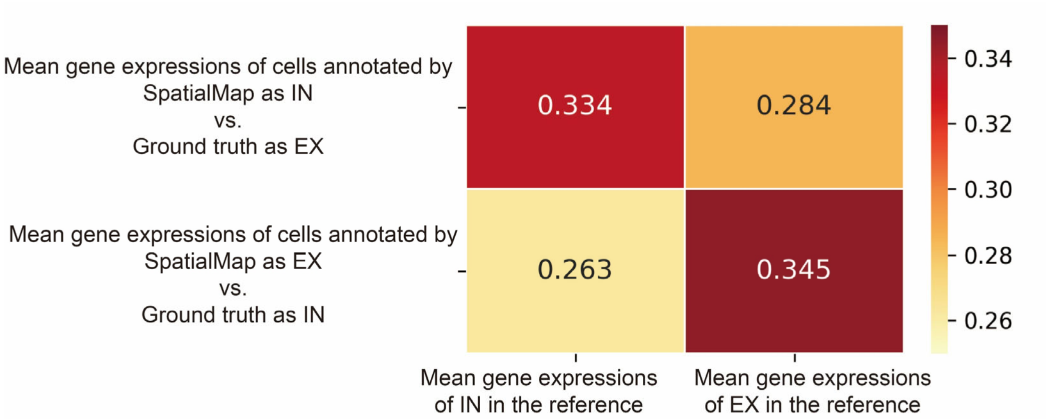


Figure S6. Cosine similarity between the conflicting cells (SpatialMap vs. Ground truth of inhibitory neurons (IN) and excitatory neurons (EX)) and reference expression profiles.

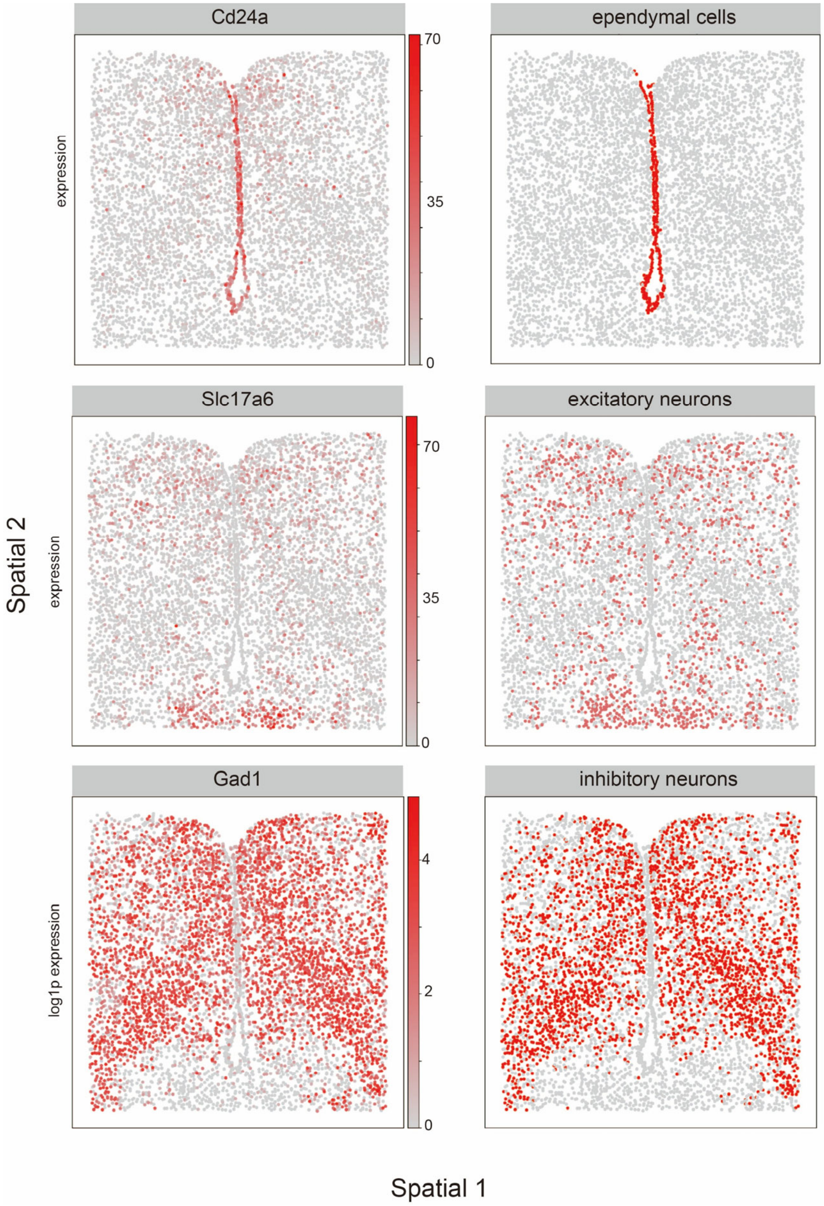


Figure S7. Spatial visualization of the mouse hypothalamic preoptic region dataset (begma=-0.04) for ependymal cells, excitatory neurons and inhibitory neurons annotated by SpatialMap with corresponding marker genes.

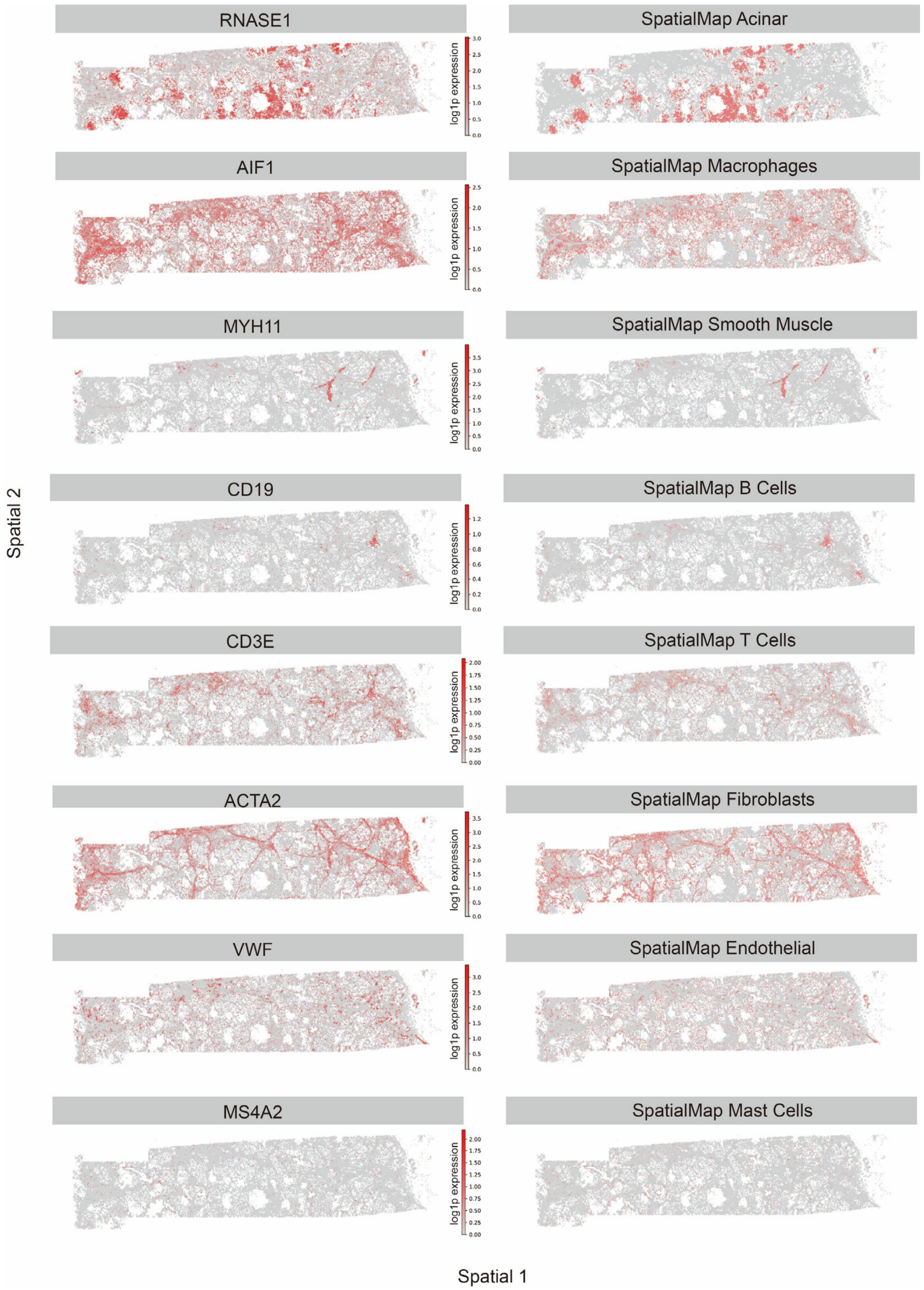


Figure S8. Spatial visualization of each cell type of SpatialMap's annotation (right) on the human pancreas dataset with corresponding marker gene (left).

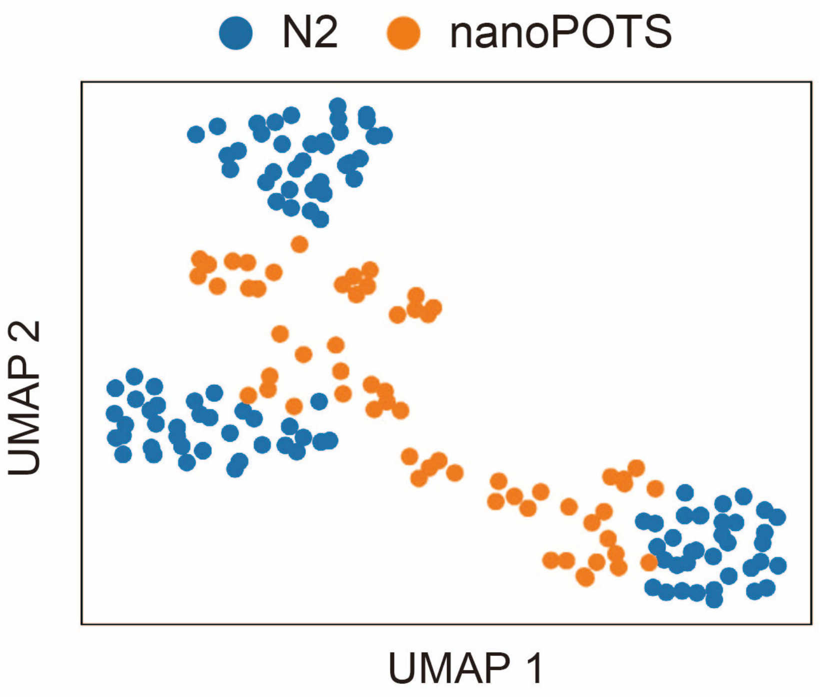

Figure S9. UMAP embeddings showing the distribution of two proteomics datasets: N2 and nanoPOTS.

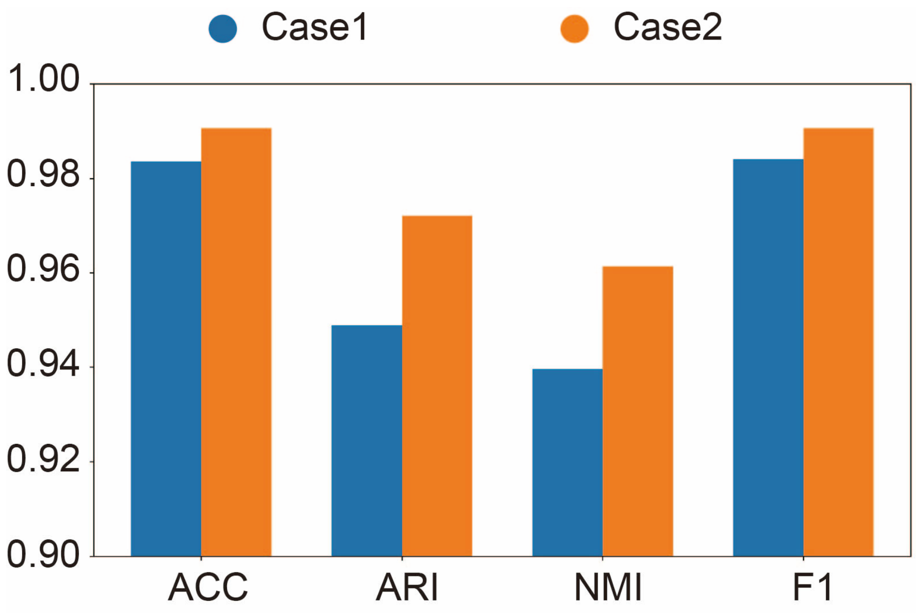


Figure S10. Quantitative metrics of SpatialMap in Case1 (N2 to nanoPOTS) and Case2 (nanoPOTS to N2) of cross-platform proteomics annotation, including ACC, ARI, NMI, F1.

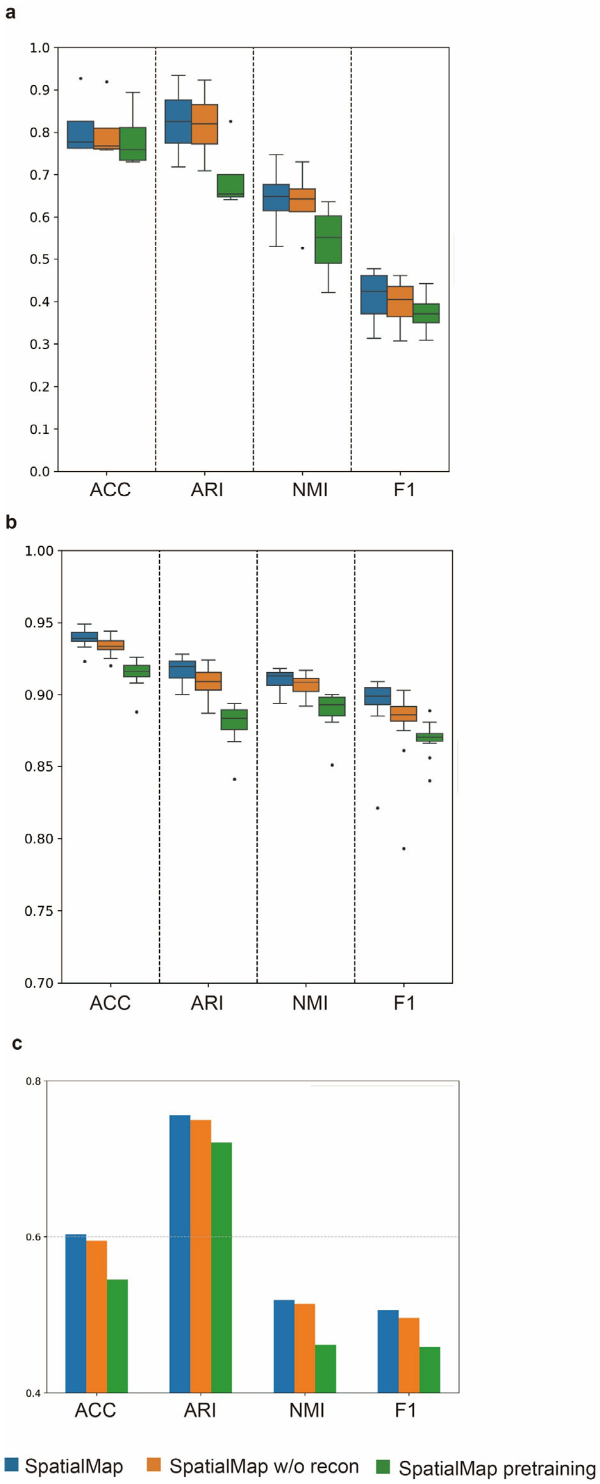


Figure S11. Ablation studies of SpatialMap's on four real-world subcellular datasets. a: human NSCLC measured by CosMx. b: mouse primary motor cortex measured by MERFISH. c: human pancreas measured by Xenium.

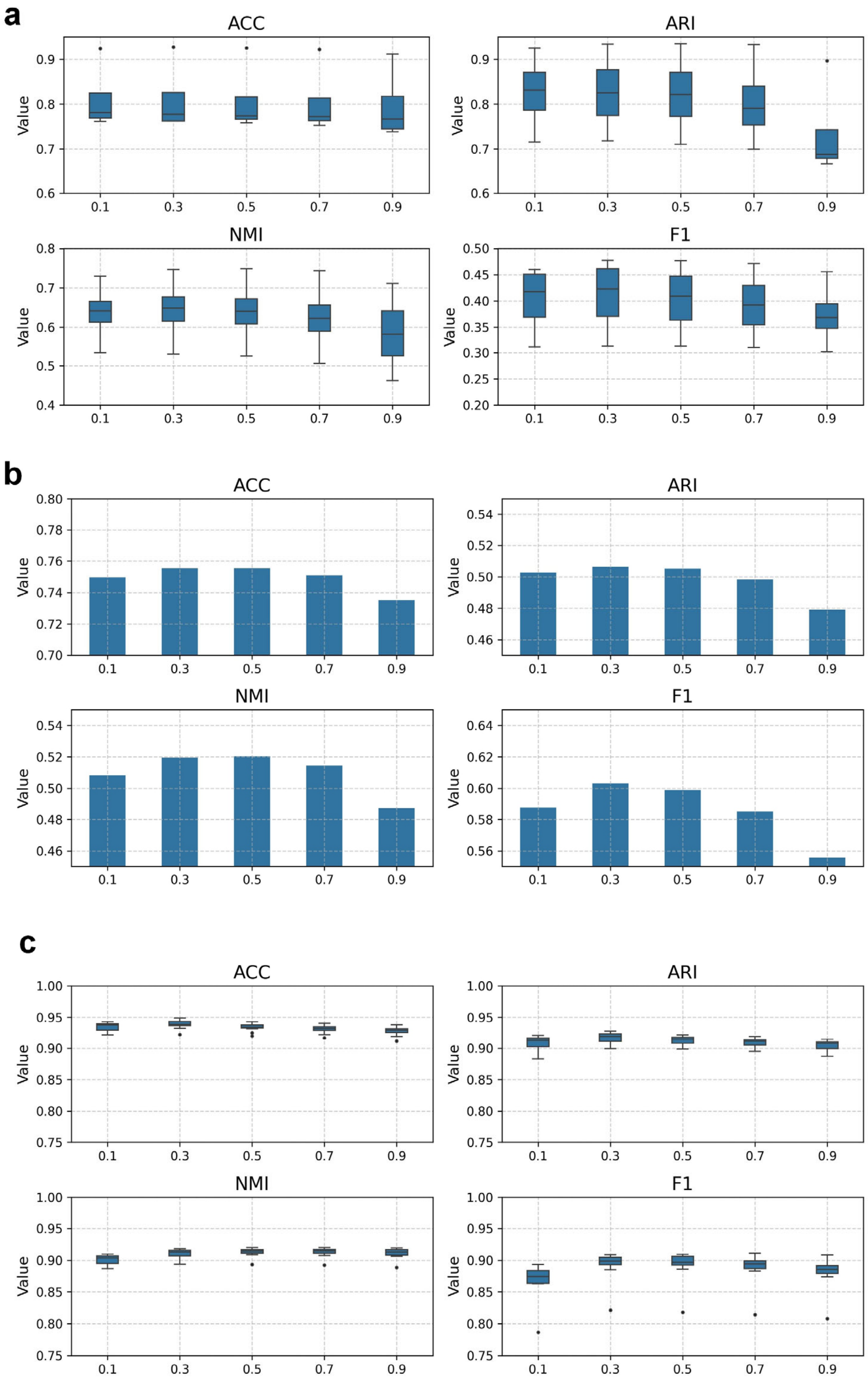


Figure S12. The influence of different β on SpatialMap's performance on three subcellular technology platform. a: the human NSCLC dataset measured by CosMx. b: the human pancreas dataset measured by Xenium. c: the mouse primary motor cortex dataset measured by MERFISH. SpatialMap adopts β=0.3 for cell type annotation.

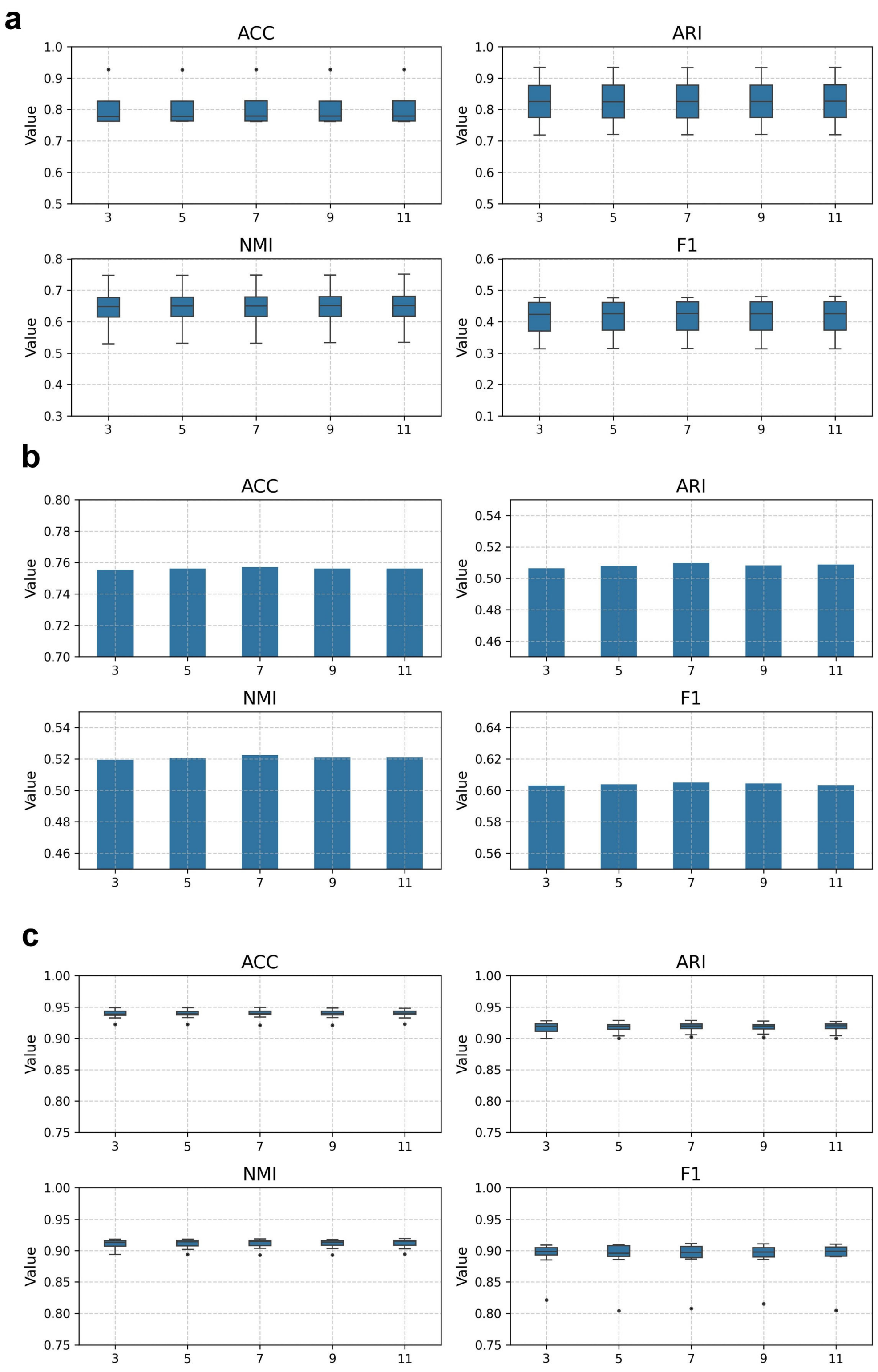


Figure S13. The influence of different k on SpatialMap's performance on three subcellular technology platform. a: the human NSCLC dataset measured by CosMx. b: the human pancreas dataset measured by Xenium. c: the mouse primary motor cortex dataset measured by MERFISH. SpatialMap shows high robustness to k.

**Table S1.** Description of all the datasets used in this study.

| Technology | Tissue | Type | Samples | Cells | Genes/Proteins | Link |
|---|---|---|---|---|---|---|
| Stereo-Seq | Mouse whole brain | Spatial transcriptomics | 1 sample from The postnatal day 7 (P7) murine whole brain sagittal section near the middle line | 96,874 | 26,497 genes | https://db.cngb.org/stomics/datasets/STDS0000139/summary |
| NanoString CosMx | Human non-small cell lung cancer (NSCLC) | Spatial transcriptomics | 4 samples: Lung-6, Lung-9-1, Lung-9-2, and Lung-13 | 419,229 | 960 genes | https://nanostring.com/products/cosmx-spatial-molecular-imager/nsclc-ffpe-dataset/ |
| scRNA-seq | Human non-small cell lung cancer (NSCLC) | Single-cell transcriptomics | / | 52,698 | 33,694 genes | https://gbiomed.kuleuven.be/english/cme/research/laboratories/54213024/scRNAseq-NSCLC |
| MERFISH | Mouse hippocampal formation | Spatial transcriptomics | 14 slices with more than 8,000 cells, including slice 19, 24, 25, 26, 27, 28, 29, 30, 31, 32, 33, 35, 36, and 37. | 262,896 | 550 genes | https://alleninstitute.github.io/abc_atlas_access/descriptions/MERFISH-C57BL6J-638850.html |
| scRNA-seq | Mouse hippocampal formation | Single-cell transcriptomics | / | 208,299 | 32,285 genes | https://alleninstitute.github.io/abc_atlas_access/descriptions/WMB-10Xv2.html |
| MERFISH | Mouse primary motor cortex (MOP) | Spatial transcriptomics | 12 samples from mouse 1 and 2 | 161,384 | 254 genes | https://doi.brainimagelibrary.org/doi/10.35077/g.21 |
| snRNA-seq | Mouse primary motor cortex (MOP) | Single-cell transcriptomics | / | 174,348 | 253 genes | https://assets.nemoarchive.org/dat-ch1nqb7 |
| MERFISH | Mouse hypothalamic preoptic | Spatial transcriptomics | 12 slices from mouse 1 | 73,655 | 155 genes | https://datadryad.org/stash/dataset/doi:10.5061/dryad.8t8s248 |
| scRNA-seq | Mouse hypothalamic preoptic | Single-cell transcriptomics | / | 31,299 | 27,998 genes | https://www.ncbi.nlm.nih.gov/geo/query/acc.cgi?acc=GSE113576 |
| 10x Genomics Xenium | Human pancreas | Spatial transcriptomics | 1 sample | 190,965 | 538 genes | https://www.10xgenomics.com/products/xenium-human-pancreatic-dataset-explorer |

| | | | | | | |
|---|---|---|---|---|---|---|
| scRNA-seq | Human pancreas | Single-cell transcriptomics | / | 47,171 | 33,538 genes | https://www.immunesinglecell.org/ |
| MERFISH | Human liver | Spatial transcriptomics | 2 samples | 1,166,496 | 550 genes | https://info.vizgen.com/ffpe-showcase |
| scRNA-seq | Human liver | Single-cell transcriptomics | T010016 and T010045 | 9,597 | 18,500 genes | http://tisch.comp-genomics.org/home/ |
| CyTOF | Human breast atlas | Single-cell proteomics | 50 samples from 38 patients | 751,970 | 34 proteins | https://data.mendeley.com/datasets/vs8m5gkyfn/1 |
| scRNA-seq | Human breast atlas | Single-cell transcriptomics | / | 52,681 | 20,437 genes | https://www.ncbi.nlm.nih.gov/geo/query/acc.cgi?acc=GSE180878 |
| N2 | Murine cell lines | Single-cell proteomics | / | 108 | 1,068 proteins | https://massive.ucsd.edu/ProteoSAFe/dataset.jsp?accession=MSV000086809 |
| nanoPOTS | Murine cell lines | Single-cell proteomics | / | 61 | 1225 proteins | https://massive.ucsd.edu/ProteoSAFe/dataset.jsp?task=09a67b73be7943739241ab540074d12f |
